# Integration of variant annotations using deep set networks boosts rare variant association genetics

**DOI:** 10.1101/2023.07.12.548506

**Authors:** Brian Clarke, Eva Holtkamp, Hakime Öztürk, Marcel Mück, Magnus Wahlberg, Kayla Meyer, Felix Munzlinger, Felix Brechtmann, Florian R. Hölzlwimmer, Julien Gagneur, Oliver Stegle

## Abstract

Rare genetic variants can strongly predispose to disease, yet accounting for rare variants in genetic analyses is statistically challenging. While rich variant annotations hold the promise to enable well-powered rare variant association tests, methods integrating variant annotations in a data-driven manner are lacking. Here, we propose DeepRVAT, a model based on set neural networks that learns burden scores from rare variants, annotations, and phenotypes. In contrast to existing methods, DeepRVAT yields a single, trait-agnostic, nonlinear gene impairment score, enabling both risk prediction and gene discovery in a unified framework. On 34 quantitative and 26 binary traits, using whole-exome-sequencing data from UK Biobank, we find that DeepRVAT offers substantial increases in gene discoveries and improved replication rates in held-out data. Moreover, we demonstrate that the integrative DeepRVAT gene impairment score greatly improves detection of individuals at high genetic risk. Finally, we show that pre-trained DeepRVAT scores generalize across traits, opening up the possibility to conduct highly computationally efficient rare variant tests.

## Introduction

The recent arrival of population-scale whole-exome and whole-genome sequencing studies^1^ vastly expands the potential to understand the genetic underpinnings of human traits. While genome-wide association studies (GWAS) on common variants have identified a compendium of trait-associated loci^2^, mapping these largely noncoding variants to affected genes as well as addressing the typically subtle effect sizes remains challenging^3,4^. In contrast, rare variants can exhibit large effects^5^, aiding the discovery of effector genes^6,7^, the unraveling of molecular mechanisms underlying traits, and in turn, the identification of potent drug targets^8–10^. Of further medical relevance, modeling of rare variant effects has recently shown promise in the identification of individuals at high risk for disease and in deriving polygenic risk scores that generalize better across populations than those based only on common variants^11^.

However, extending the GWAS strategy to rare variants must contend with large numbers of low-frequency variants, leading to low statistical power due to sparsity and an increased multiple testing burden. To compensate, rare variant association testing (RVAT) methods aggregate rare variants at the level of genomic regions, typically genes^12,13^. Furthermore, the presence or magnitude of impact on gene function for a given rare variant cannot typically be inferred directly. Therefore, RVAT methods rely on functional predictions of variant effect^12,14–16^, such as conservation scores or variant effect predictions for splicing, gene expression or protein structure^17–19^ to prioritize putatively impactful variants.

Burden testing, a common RVAT strategy, uses annotations to filter presumably uninformative variants and to weight informative ones. These weights are then aggregated to a gene-level burden score and tested for association with discrete or quantitative traits^20–25^. Complementary to burden tests, variance component tests, which can account for both protective and deleterious variants, also use annotations for filtering and weighting variants as part of a kernel function^13,26,27^. Recently proposed RVAT methods based on variance component tests convincingly demonstrated the added value of incorporating a broad spectrum of annotations, either by conducting a meta-analysis over different test types and annotations^28^, or using specialized kernels tailored to different annotation types^29^ (**Methods**; **Supp. Table 1**). However, like earlier methods, *ad hoc* variant filtering and weighting schemes remain integral components of existing workflows. The few RVAT methods that infer annotation weights from data and integrate multiple annotations are computationally prohibitively demanding, and in practice limited in the number of annotations and in the flexibility of the scoring function that can be considered^30,31^. Finally, because of these limitations none of these methods lends itself to phenotype prediction, thus greatly limiting their utility for applications in personalized medicine (**Methods**; **Supp. Table 1**).

To address these issues, we present DeepRVAT (Deep Rare Variant Association Testing). Our framework uses a deep set network for modeling traits by integration of rare variant annotations. The model handles variable numbers of rare variants per individual, leverages dozens of continuous or discrete annotations, and accounts for both additive and interaction effects. All model parameters are learned directly from the training data, minimizing the need for *ad hoc* modeling choices that characterize existing methods. The trained DeepRVAT model gives rise to a single, trait-agnostic gene burden scoring function. This offers several key advantages over variance-component tests. Firstly, it can be used for different genetic analyses, including rare variant association tests and to refine polygenic risk prediction to account for rare variant effects. Second, it retains calibration when testing for association with imbalanced binary traits, which was problematic for alternative methods using rich annotations. Finally, DeepRVAT provides large gains in computational efficiency over alternative methods. We provide pre-trained DeepRVAT models for direct application on new data sets, as well as a pipeline for retraining the model from scratch, for example to incorporate additional annotations.

We validate the model using simulations before applying it to 34 quantitative and 26 binary traits, using whole-exome-sequencing data from 167,245 UK Biobank (UKBB) individuals, demonstrating a larger number of discoveries compared to existing methods. Additionally, we demonstrate improved replication of DeepRVAT associations in held-out individuals from UKBB. Finally, we combine DeepRVAT gene impairment scores with polygenic risk models, which yields improved prediction accuracy for extreme value phenotypes in UKBB traits.

## Results

### A deep set network-based RVAT framework

DeepRVAT is an end-to-end model (**Fig. 1a**) that first accounts for nonlinear effects from rare variants on gene function (gene impairment module) to then model variation in one or multiple traits as linear functions of the estimated gene impairment scores (phenotype module). The gene impairment module (**Fig. 1b**) outputs a gene- and trait-agnostic gene impairment score estimating the combined effect of rare variants, thereby allowing the model to generalize to new traits and genes. Technically, a deep set neural network^32^ architecture is employed to aggregate the effects from multiple discrete and continuous annotations for an arbitrary number of rare variants. This architecture captures additive and nonlinear interactions and does not rely on *a priori* assumptions about the relevance of individual annotations, such as common assumptions about the relationship between allele frequency and variant effects^20,22^ (**Methods**).

**Figure 1.**
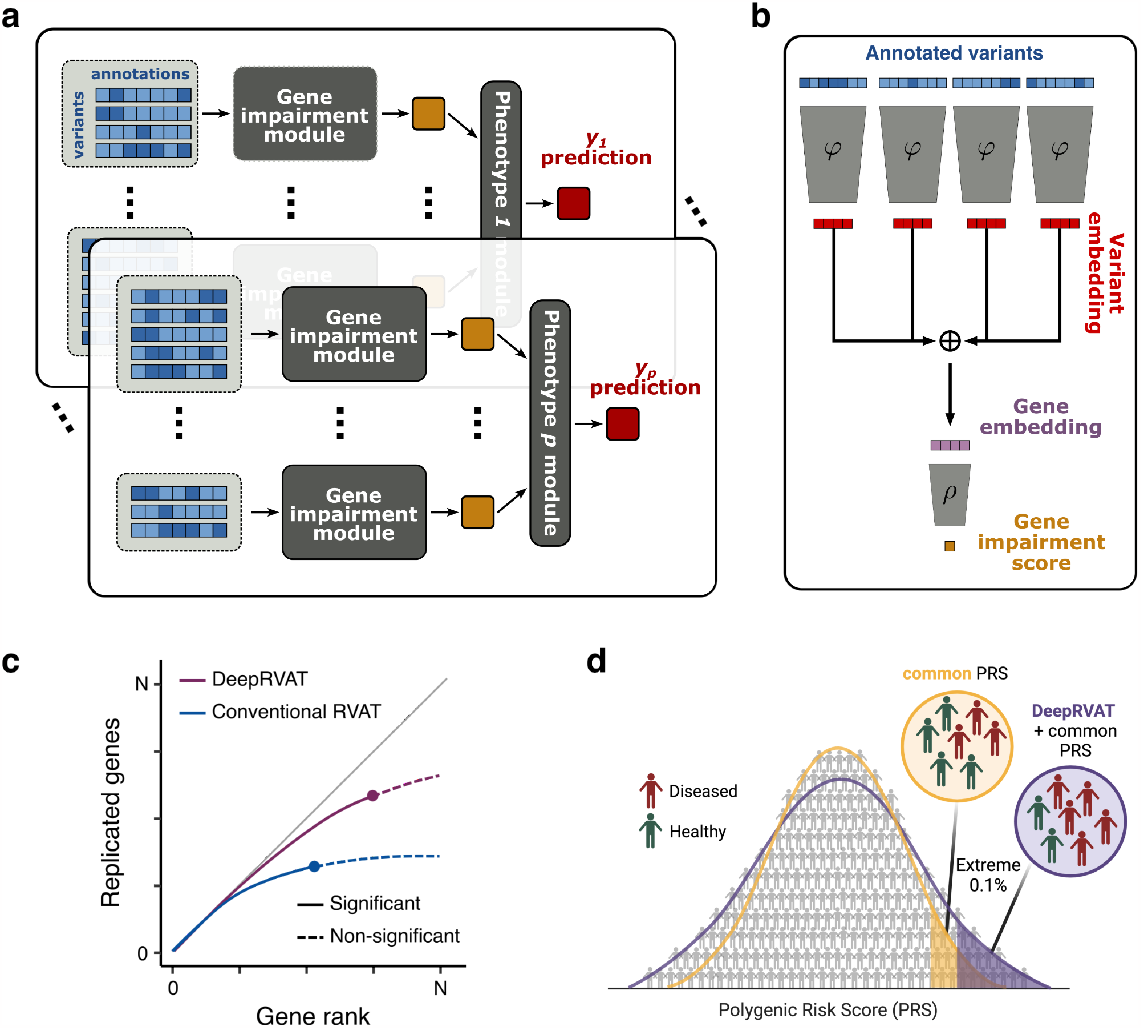
DeepRVAT model overview and downstream use cases. **(a)** DeepRVAT predicts phenotype from genotype by modeling gene impairment from individual-level rare variants and annotations (Gene impairment module, **b**), and additive trait-specific gene effects. **(b)** DeepRVAT gene impairment module. Gene impairment scores are estimated from personalized rare variant profiles for each individual and gene combined with an arbitrary number of variant annotations as input. Variants are fed through an embedding network, *φ*, to compute a variant embedding, followed by permutation-invariant aggregation to yield a gene embedding. A second network, *ρ*, estimates the gene impairment score. All parameters of the gene impairment module are shared across genes and traits. **(c,d)** Downstream analyses based on the DeepRVAT impairment score. **(c)** Gene discovery. The trained DeepRVAT impairment module can be used as a scoring function to define association tests for gene discovery, which are assessed for number of discoveries and replication in held-out data. **(d)** Polygenic risk prediction incorporating rare variants. The DeepRVAT gene impairment module can be used as input to train predictive models for phenotype from genotype, for example, to improve risk stratification based on polygenic risk scores (PRS) with common variants. Created with BioRender.com
.

For each trait used in model training, a set of informative seed genes that are associated with the given trait are required. The DeepRVAT software offers an integrated workflow that employs conventional rare-variant association tests to identify seed genes (**Methods**). Next, the gene impairment module is trained end-to-end together with the phenotype modules on the seed genes. Once trained, the DeepRVAT gene impairment module can be applied to gene-trait combinations not used during training to test for genetic associations using the established principles of conventional burden tests (**Fig. 1c**). The estimated gene impairment scores from the trained DeepRVAT module can also be used to predict phenotype from genotype, thereby providing a flexible way to derive polygenic risk scores^11,33^ that account for rare variant effects (**Fig. 1d)**. The training time of DeepRVAT scales linearly with the number of individuals, and association testing is highly efficient (**Supp. Fig. 1.1**), thereby enabling applications to phenome-wide association studies on biobank-scale datasets.

### Model validation using simulated data

To validate DeepRVAT for gene discovery, we considered a semi-synthetic dataset derived from whole exome sequencing (WES) from 167,245 caucasian UK Biobank individuals (November 2020, 200k WES release^34^; **Methods**). We annotated all 12,704,497 WES variants using minor allele frequency (MAF), VEP^35^ consequences, missense variant impact scores (SIFT^14^ and PolyPhen2^16^), and omnibus statistical deleteriousness scores (CADD^15^ and ConDel^36^), as well as predicted annotations for effects on protein structure (PrimateAI^18^), and aberrant splicing (AbSplice^17^). To create synthetic phenotypic data, we first simulated impairment scores for genes designated to be causally associated to the trait. To this end, we stochastically assigned each variant an effect based on its annotations (**Supp. Fig. 2.1, Methods**), followed by aggregation for each gene. Finally, the individual phenotype values were simulated as a linear function of the gene impairment scores and additive effects from covariates mimicking age and sex, and independent Gaussian noise.

After confirming statistical calibration (**Fig. 2a, Supp. Fig. 2.2**), we compared the power of DeepRVAT to conventional burden testing as well as variance component testing with SKAT^26^, following Karczewski et al.^22^, i.e., considering either predicted loss of function (pLOF) or missense variants and with weighting based on minor allele frequency (MAF; **Methods**). Notably, existing methods rely on a predefined MAF cutoff. To investigate the impact of a possible misspecification of the MAF cutoff, we varied the frequencies of the simulated causal variants (**Fig. 2b**, top to bottom, **Methods**). For each simulation setting, we applied the considered methods for alternative cutoff values of the MAF frequency (<0.01%, <0.1%, <1%; **Fig. 2b**, left to right). Whereas the power of conventional tests was greatly affected by the alignment of the simulated MAF distribution of causal variants and the MAF filter, DeepRVAT was consistently well powered, including in settings for which additional non-causal variants were incorporated in the MAF cutoff. Collectively, these results support that DeepRVAT is able to estimate an appropriate weighting for variants from the data, including implicit prioritization of variants by MAF.

**Figure 2.**
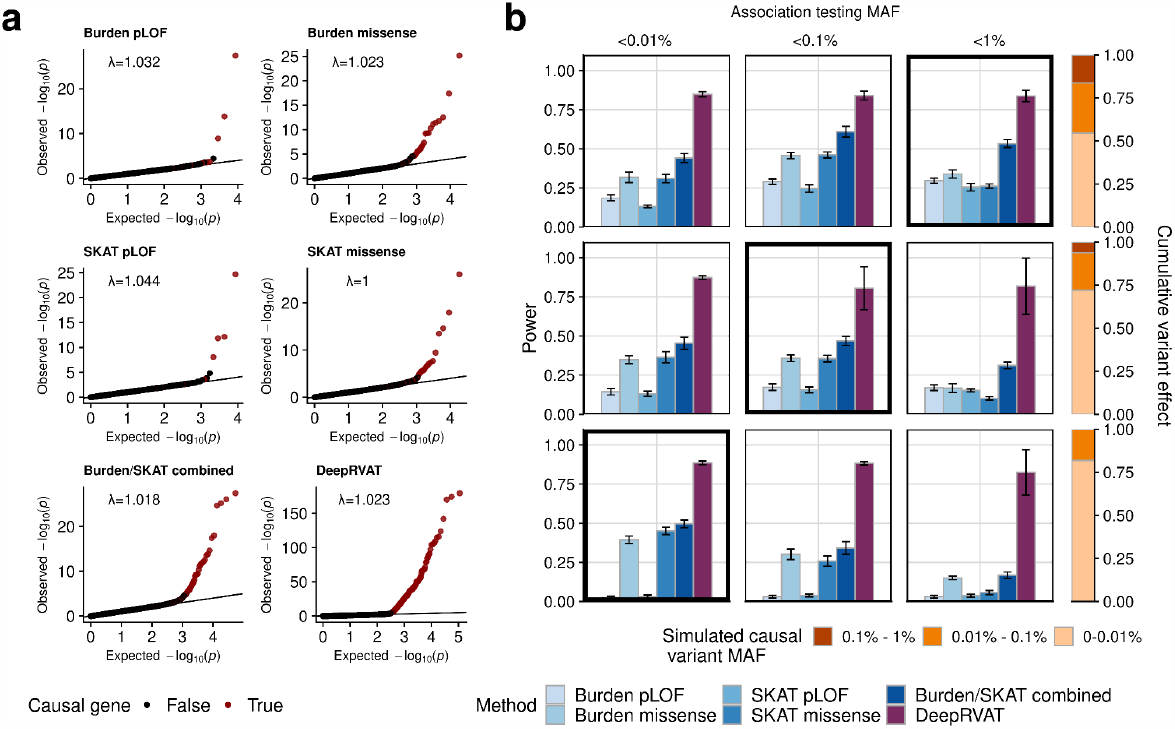
Method assessment using simulated data. Considered were DeepRVAT using complete annotations, conventional burden tests and SKAT, each based on either pLOF or missense variants, and the combination of these four methods (Burden/SKAT combined). **(a)** Assessment of calibration for DeepRVAT and alternative methods. Shown are Q-Q plots of the expected versus the observed *p*-value distribution. Lambda denotes the genomic inflation factor. Genes simulated to be causal in red; non-causal genes in black. **(b)** Power comparison for alternative MAF distributions of simulated causal variants (rows) as well as MAF filters applied to determine input variants for each of the considered tests (columns). Shown is average power (bar height) and standard deviation (error bars) across 10 simulation runs. Results on the diagonal correspond to the setting for which the MAF filter of the RVAT tests are aligned to the simulation setting; others correspond to varying degrees of MAF filter misspecification. Stacked bar plots on the right for each row denote the relative contribution of variants in different frequency bins for the respective simulation settings (averaged across all simulation runs).

Similarly, we considered altering the proportion of rare variants that are selected to contribute to gene impairment effects (**Supp. Fig. 2.1, Supp. Fig. 2.3a**) and a rank-based evaluation criterion (**Supp. Fig. 2.3b**). Across these settings, DeepRVAT was consistently better powered than alternative methods. In sum, the benchmark using simulated data demonstrates that DeepRVAT yields results that are robust to a range of key parameters, including the MAF spectrum and the proportion of causal variants.

### DeepRVAT recovers a larger number of gene-trait associations using rare variants in UK Biobank

Next, we applied DeepRVAT to rare variants (MAF < 0.1%) from whole-exome data from the UK Biobank, considering the same 167,245 caucasian individuals as used in the simulation study. We used a total of 33 variant annotations (**Supp. Table 5; Supp. Fig. 3.1**), expanding the set in the simulation study by variant effect predictions for epigenetic markers in cell types from ENCODE^37^ and the Roadmap Epigenomics^38^ projects (using principal components of DeepSEA^19^ predictions; **Supp. Fig. 3.2; Methods**), as well as binding to 6 RNA-binding proteins (selected predictions from DeepRiPe^39^; **Methods**).

We used 21 quantitative traits of various categories (Supp. Table 3; Methods) to train DeepRVAT, followed by genome-wide association testing. Across all traits, DeepRVAT identified 516 gene-trait associations (FDR < 5%; Supp. Table 6), which corresponds to a 105% increase compared to a widely used baseline approach combining burden and SKAT, a 16% increase compared to Monti et al. and a similar number of associations as retrieved by STAAR (**Fig. 3a**). However, whereas DeepRVAT retained statistically calibrated p-vaues, STAAR suffered from poorly calibrated test statistics (**Supp. Fig. 3.3**), thus precluding a meaningful comparison with this approach. At the level of individual traits, DeepRVAT identified the largest number of associations among all considered methods (excluding STAAR) for 14 out of the 21 traits (8 out 21 traits when including STAAR).

**Figure 3.**
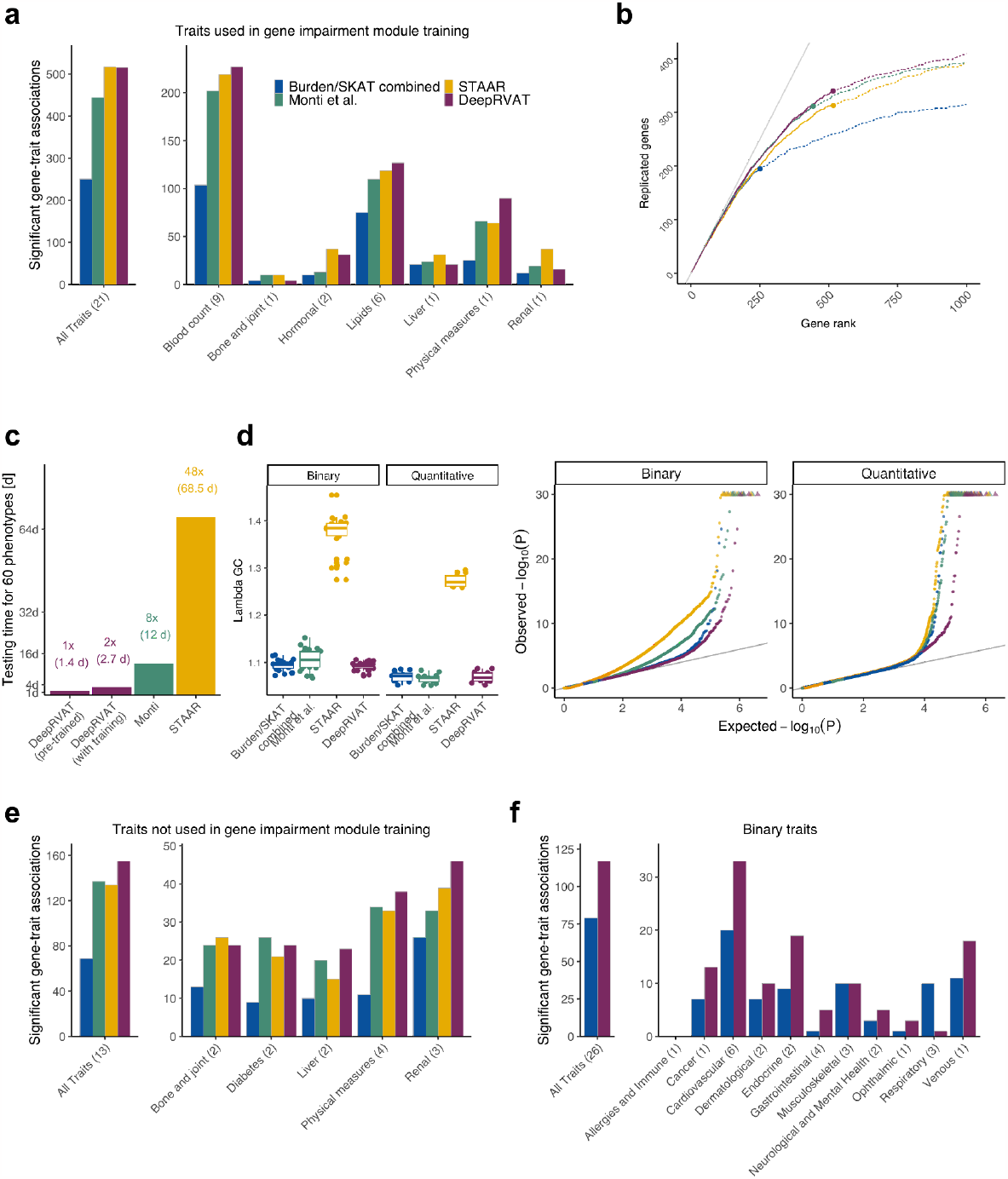
Application of DeepRVAT to gene discovery using WES and 60 traits from 167,245 UK Biobank individuals. Considered are DeepRVAT, the combination of burden and SKAT tests, each on both missense and pLOF variants (Burden/SKAT combined), STAAR, and the approach from Monti et al. **(a)** Application to 21 quantitative traits used for training DeepRVAT. Left: Cumulative number of gene-trait associations discovered by alternative methods (FDR < 5%). Right: Number of gene-traits associations broken down by trait category. **(b)** Replication of the cumulative discoveries across traits by alternative methods as in **a** in a larger cohort (cohort supersets; UK Biobank full release^20,22^, **Supp. Table 7**). Shown is, for each method, the total number of gene-trait associations that were also discovered in the larger cohort (according to the methodology of the respective studies) as a function of the rank of their nominal significance. Circles indicate the rank position that corresponds to FDR < 5% (as in **a**). The gray line corresponds to a replication rate of one. **(c)** Estimated computation when testing for association of 20,000 genes across 60 phenotypes. Numbers denote multiples of the compute time for the pre-trained DeepRVAT model. Shown are empirical runtimes on a workstation with an AMD Ryzen Threadripper PRO 5975WX CPU and an NVIDIA RTX 4090 24GB GPU. **(d)** Calibration of DeepRVAT and alternative methods on quantitative and binary traits not considered during DeepRVAT training. Left: Distribution of the genomic inflation factor across traits. Right: Q-Q plots of expected vs. observed *p*-values. **(e)** Application to 13 quantitative traits not considered during DeepRVAT training. Left and right analogous to **a**. **(f)** Application of DeepRVAT to 26 binary traits. Left: Cumulative number of gene-trait associations discovered by DeepRVAT and the “Burden/SKAT combined” method (FDR < 5%). Right: Number of gene-trait associations from left plot broken down by trait. Monti et al.’s method and STAAR are not shown given their poorly calibrated test statistics and infliated discoveries (c.f. **Fig. 3c, Supp. Fig. 3.3b,c**).

Next, we evaluated the validity of the discoveries by assessing their replication in at least one of two studies on the full UK Biobank WES cohort (N=454,787 and 394,841 individuals), which employed analysis strategies based on SKAT and burden testing. Notably, across a wide range of nominal significance ranks, the replication rate of DeepRVAT exceeded the replication of discoveries of alternative methods (**Fig. 3b**). This suggests that not only does DeepRVAT have an improved capacity to detect subtle associations compared to existing methods, but it is also less susceptible to spurious ones.

Once DeepRVAT is trained, the gene impairment module can be applied independently of the training traits, allowing for highly efficient association analysis (**Fig. 3c, Supp. Fig. 1.1**). To assess the universality of the DeepRVAT impairment scoring, we extended our investigation to 13 additional quantitative and 26 binary traits that were not considered during training. In contrast to Monti et al. and STAAR, where we find poor calibration for binary traits, in concordance with other studies^40^, DeepRVAT retained robust calibration also for binary traits (**Fig. 3d, Supp. Fig. 3.3**). Furthermore, DeepRVAT again yielded an increased number of discoveries compared to alternative methods. The model identified 155 gene-trait associations (FDR < 5%) for quantitative traits, representing a 124% increase compared to the combination of burden and SKAT (**Fig. 3e**), and 117 discoveries for the binary traits, which corresponds to a 48% increase compared to the only calibrated alternative methods (**Fig. 3f**).

Finally, we considered DeepRVAT with a reduced set of annotations (MAF, pLOF and missense status), as well as a simplified model architecture with a fully linear gene impairment module. These models yielded lower performance than the full DeepRVAT model when considering discoveries and replication (**Supp. Fig. 3.4**), but still significantly outperformed baseline models. This indicates that the data-driven scoring function, the ability to consider a larger number of annotations, and the capacity to capture nonlinear effects all jointly contribute to the overall performance. This is further supported by an analysis of the relevance of different annotations in DeepRVAT (**Supp. Fig. 3.5**), which suggests that non-canonical annotations such as DeepSEA predictions add value even for coding variants.

### DeepRVAT improves phenotype prediction by integrating rare variants

An important advantage of DeepRVAT over approaches that combine different annotations by aggregating across multiple tests such as STAAR and Monti et al. is that the gene impairment scores can be used for phenotype predictions. To assess the added value of DeepRVAT for this task, we trained regression models that combine polygenic risk scores (PRS) derived from common variants^33^ with gene impairment scores from DeepRVAT. For comparison, we trained analogous models in which the gene impairment was based on single-variant annotations as in conventional gene burden tests. All models were evaluated using 5-fold cross-validation across individuals. For each fold, conventional RVAT methods (*Burden/SKAT combined*; previous section) or DeepRVAT were employed for gene discovery (FDR < 0.05) followed by training of the phenotype prediction model on the same training set (**Methods**). All models were evaluated on the held-out validation set.

We first applied this strategy to a regression task to predict the same quantitative traits as in the previous section (**Fig. 4a**). (One trait was excluded due to not having an available PRS.) Compared to models based on conventional RVAT burdens, the relative prediction improvement of DeepRVAT vs. common variant PRS was significantly larger across these 33 traits (maximum improvement of 8.04% for Alkaline phosphatase). However, the overall contribution of rare variants was modest (median improvement of 0.86% for DeepRVAT), which is consistent with the expectation that common variants explain most of the trait variation across the population. We hypothesized, however, that the importance of rare variants would be higher when applying the same modeling strategy to predicting the individuals in the bottom or top 1% of the phenotypic range (**Fig. 4b**). Indeed, the relative improvement for predicting the extreme percentiles was considerably larger (maximum improvement using DeepRVAT of 233.02% for predicting individuals with low Alkaline phosphatase measurement), and we again found significant advantages of the DeepRVAT impairment score compared to alternative burden measures. Dissection analyses, combining alternative methods for gene discovery and burden score estimation, showed that the gene impairment score and the set of identified genes individually contribute to the added value of DeepRVAT (**Supp. Fig. 4.1**).

**Figure 4.**
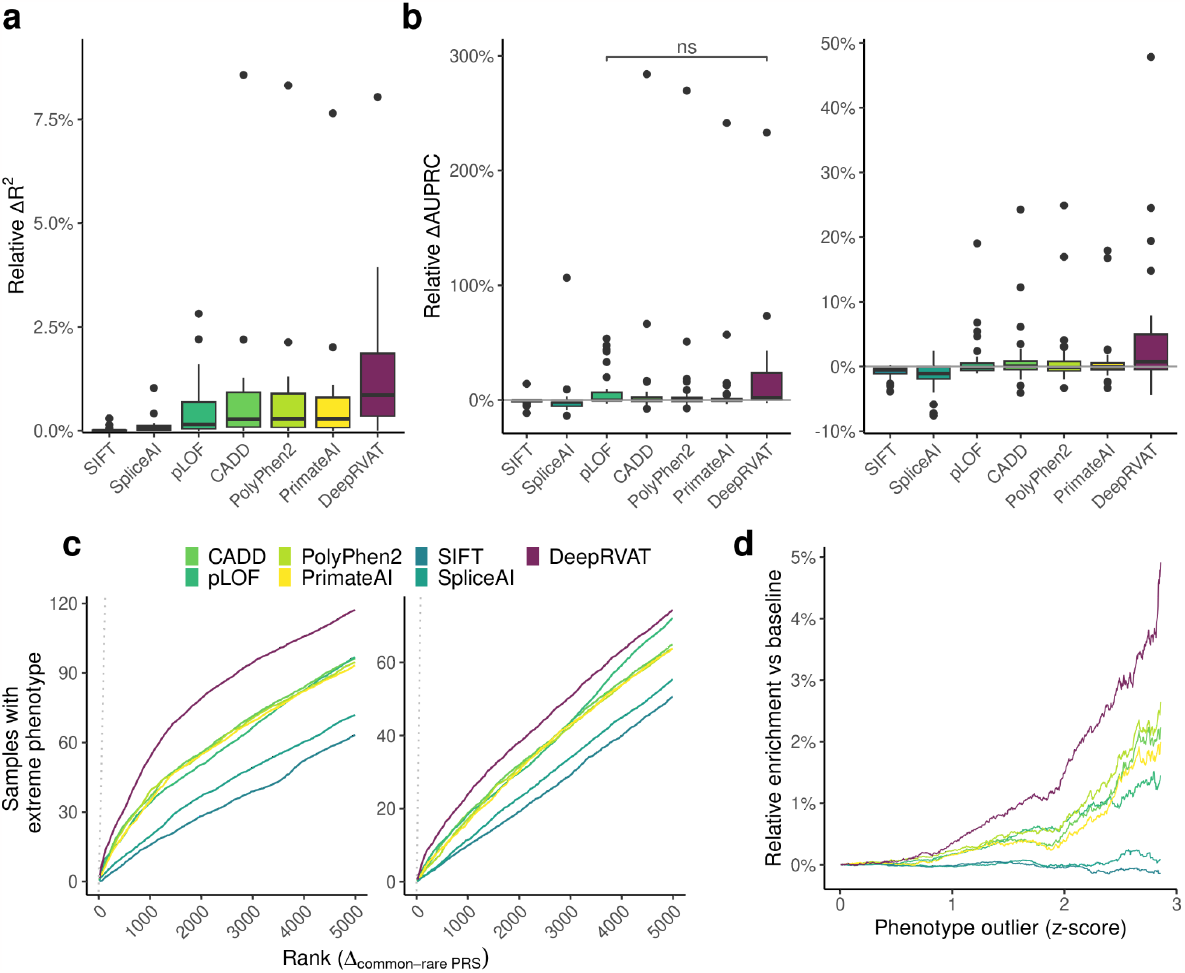
Application of DeepRVAT and alternative gene burdens for phenotype prediction. Assessment of alternative rare variant gene burdens as features for phenotype prediction of 33 UK Biobank traits. Considered were polygenic risk scores derived from common variants, combined with either DeepRVAT or one of six alternative rare burden measures, evaluated by 5-fold cross-validation. **(a)** Relative improvement of the prediction performance when including rare variant gene burdens in a linear regression model based on common variant PRS. Shown are relative differences in the coefficient of determination (R^2^) for models that account for rare variants versus a common-variant PRS model. **(b)** Analogous comparison as in **a**, however considering a logistic regression model to stratify individuals in the bottom or top 1% of the phenotypic distribution. Shown are relative differences in the area under the precision-recall curve (AUPRC) between a model that accounts for rare variants versus a common variant PRS model. Unless indicated otherwise (“ns”), the relative gains of including the DeepRVAT burden compared to alternative methods as in **a** and **b** are significant (p < 0.05, one-sided Wilcoxon test). **(c)** Rank-based enrichment of individuals with extreme value phenotypes (top or bottom 1%) among individuals with strongly deviating predictions when using a model that combines PRS and a rare variant burden. Shown is the number of individuals with extreme-value phenotypes when ranking individuals by the magnitude of deviation using a model that accounts for rare variants versus a common-variant PRS model. Shown are average enrichment values across all 33 traits. **(d)** Enrichment of outlier phenotype predictions (exceeding the 99% quantile) in individuals with extreme phenotypes. Shown is the enrichment of outlier individuals (defined as exceeding a certain Z-score cutoff, x-axis) for models that account for rare variants relative to the enrichment obtained from a common-variant PRS model. Shown are average enrichment values across all 33 traits.

Next, we focused on the set of individuals with the strongest deviations between the rare variant model and the common PRS predictor. Particularly with DeepRVAT, individuals showing the greatest deviation in predicted phenotypic values were more likely to exhibit extreme phenotype values (bottom or top 1%; **Fig. 4c**). Similarly, individuals with outlying phenotype predictions from the DeepRVAT model were more prominently enriched for extreme phenotype values compared to alternative rare variant predictors (**Fig. 4d**). At the most extreme phenotype threshold (z-score ≥ 2.86), DeepRVAT showed a 4.9% greater enrichment of phenotypic outliers in its 99th percentile predictions compared to the common variant PRS model, outperforming the most competitive rare variant method, PolyPhen2, which only gave a 2.6% improvement. These findings demonstrate the benefits of DeepRVAT gene impairment scores for phenotype prediction, particularly in identifying individuals with outlier phenotypes, compared to conventional PRS based on common variants and alternative rare variant models.

## Discussion

We have introduced DeepRVAT, a data-driven deep-learning model for association testing and phenotype prediction based on rare variants. Unlike existing methods, DeepRVAT infers the relevance of different annotations and their combination directly from data. In so doing, DeepRVAT eliminates the need for *post hoc* aggregation of test results derived from individual annotations using multiple testing schemes. DeepRVAT combines the benefits of a flexible deep neural network architecture with a calibrated statistical testing framework. It leverages the flexibility of deep neural networks to non-linearly integrate rare variant annotations while still offering a calibrated statistical framework for gene-phenotype association testing. DeepRVAT significantly outperformed state-of-the-art methods in gene discovery for 60 traits from the UK Biobank, leading to a substantial increase in performance and retrieving gene-trait associations with higher replication rates in held-out data.

In addition to improving the discovery of gene-trait associations over alternative methods utilizing rich variant annotations, DeepRVAT represents a conceptual advance by estimating a universal gene impairment score. We have demonstrated the utility of this impairment score for rapid gene-trait association testing by considering traits that were not seen by the model during training. For binary traits in particular, DeepRVAT offers improved calibration compared to alternative methods that leverage rich annotations, especially STAAR^40^. A second opportunity provided by the impairment score is to estimate genetic predisposition of individuals by accounting for variants from the full frequency spectrum. We have demonstrated this by combining DeepRVAT with polygenic risk scores based on common variants, finding considerable benefits, particularly for extreme phenotypes, and outperforming conventional burden scores based on single annotations. DeepRVAT is provided as a user-friendly software package that supports both *de novo* training of gene impairment modules as well as application of pretrained ones, each with substantial improvements in computational efficiency over existing methods.

Although we found that DeepRVAT advances the state-of-the-art in two use cases central to genetics, the model is not free of limitations. First, choosing the most relevant annotations is a challenge for all annotation-based methods. DeepRVAT is no exception, yet it can cope with a potentially very large number of annotations by learning how to weight them directly from the data. Second, we benchmarked DeepRVAT using exome sequencing data, but whole genome sequencing promises valuable insights into non-coding regions with numerous rare variants of uncertain impact. We hypothesize that the added value of DeepRVAT, which incorporates rich variant annotations, might be even larger in this context. Finally, while we showed that DeepRVAT is robust to various rare variant frequency cutoffs, it is still based on a dichotomy between rare and common variants. Recent reports have shown substantial overlap between GWAS loci and rare-variant associations^10,11,20^. These insights suggest the potential for future developments to jointly model and estimate rare and common variant effects in a unified framework.

The integration of rare variants will remain a major topic in quantitative genetics modeling. Among rare variant association methods, DeepRVAT belongs to the class of approaches that model gene impairment, a strategy that underpins high-impact variant filters in burden tests^20,22,23^ or protein function impairment^25^. Our contribution to this family of approaches is to define a better-optimized impairment score by allowing the integration of different annotations and learning a more flexible model directly from cohort data. Importantly, we found this gene impairment score to generalize well across traits. This transferability, combined with DeepRVAT’s computational efficiency, is an important feature that will facilitate its application to conduct rapid rare-variant phenome-wide association studies (PheWAS), to perform rare variant associations in smaller cohorts from case-control studies, or to analyze cohorts from rare disease and cancer.

## Acknowledgements

The authors wish to thank Marc Jan Bonder, Manu Saraswat, Andrea Senacheribbe, Kai Ueltzhöffer, Danai Vagiaki, and Hana Susak for helpful discussions. This research has been conducted using data from UK Biobank, a major biomedical database (project IDs 25214, 44108, and 81358). B.C., K.M, M.M., F.M., & M.W. were supported through state funds approved by the State Parliament of Baden-Württemberg for the Innovation Campus Health + Life Science Alliance Heidelberg Mannheim. E.H. was funded in part by the Helmholtz Association under the joint research school Munich School for Data Science - MUDS.

## Author contributions

B.C., E.H., J.G., & O.S. conceived the method. F.B. made additional conceptual contributions to the method. B.C., E.H., H.O., & F.H. prepared the data. B.C., E.H., H.O., K.M, M.M., F.M., & M.W. implemented the methods and analyzed the data. B.C., E.H., J.G., & O.S. interpreted the results and wrote the paper.

## Methods and data

A complete methods document with all relevant details is provided as *Supplementary methods*.

## Supplementary Data

Supplementary methods: SupplementaryMethods.pdf

Supplementary figures and table captions: SupplementaryInformation.pdf Supplementary tables: SupplementaryTables.xlsx

## Code availability

The code to run DeepRVAT can be found here: https://github.com/PMBio/deeprvat/

The code to run all analysis done in this paper and re-generate the figures can be found here: https://github.com/PMBio/deeprvat-analysis/

## Data availability

The genetic, phenotype, and covariate data are protected and are only available to researchers who have valid and approved research applications for these data within the UK Biobank (www.ukbiobank.ac.uk).

## References

1. Bycroft, C. et al. The UK Biobank resource with deep phenotyping and genomic data. Nature 562, 203–209 (2018).

2. Buniello, A. et al. The NHGRI-EBI GWAS Catalog of published genome-wide association studies, targeted arrays and summary statistics 2019. Nucleic Acids Res. 47, D1005–D1012 (2019).

3. Claussnitzer, M. et al. A brief history of human disease genetics. Nature 577, 179–189 (2020).

4. Tam, V. et al. Benefits and limitations of genome-wide association studies. Nat. Rev. Genet. 20, 467–484 (2019).

5. Bomba, L., Walter, K. & Soranzo, N. The impact of rare and low-frequency genetic variants in common disease. Genome Biol. 18, 77 (2017).

6. Sazonovs, A. et al. Large-scale sequencing identifies multiple genes and rare variants associated with Crohn’s disease susceptibility. Nat. Genet. 54, 1275–1283 (2022).

7. Gao, X. R., Chiariglione, M. & Arch, A. J. Whole-exome sequencing study identifies rare variants and genes associated with intraocular pressure and glaucoma. Nat. Commun. 13, 7376 (2022).

8. Momozawa, Y. & Mizukami, K. Unique roles of rare variants in the genetics of complex diseases in humans. J. Hum. Genet. 66, 11–23 (2021).

9. Nelson, M. R. et al. The support of human genetic evidence for approved drug indications. Nat. Genet. 47, 856–860 (2015).

10. Weiner, D. J. et al. Polygenic architecture of rare coding variation across 394,783 exomes. Nature 614, 492–499 (2023).

11. Fiziev, P. P. et al. Rare penetrant mutations confer severe risk of common diseases. Science 380, eabo1131 (2023).

12. Povysil, G. et al. Rare-variant collapsing analyses for complex traits: guidelines and applications. Nat. Rev. Genet. 20, 747–759 (2019).

13. Lee, S., Abecasis, G. R., Boehnke, M. & Lin, X. Rare-Variant Association Analysis: Study Designs and Statistical Tests. Am. J. Hum. Genet. 95, 5–23 (2014).

14. Kumar, P., Henikoff, S. & Ng, P. C. Predicting the effects of coding non-synonymous variants on protein function using the SIFT algorithm. Nat. Protoc. 4, 1073–1081 (2009).

15. Rentzsch, P., Witten, D., Cooper, G. M., Shendure, J. & Kircher, M. CADD: predicting the deleteriousness of variants throughout the human genome. Nucleic Acids Res. 47, D886–D894 (2019).

16. Adzhubei, I. A. et al. A method and server for predicting damaging missense mutations. Nat. Methods 7, 248–249 (2010).

17. Wagner, N. et al. Aberrant splicing prediction across human tissues. Nat. Genet. 55, 861–870 (2023).

18. Sundaram, L. et al. Predicting the clinical impact of human mutation with deep neural networks. Nat. Genet. 50, 1161–1170 (2018).

19. Zhou, J. & Troyanskaya, O. G. Predicting effects of noncoding variants with deep learning–based sequence model. Nat. Methods 12, 931–934 (2015).

20. Backman, J. D. et al. Exome sequencing and analysis of 454,787 UK Biobank participants. Nature 599, 628–634 (2021).

21. Jurgens, S. J. et al. Analysis of rare genetic variation underlying cardiometabolic diseases and traits among 200,000 individuals in the UK Biobank. Nat. Genet. 54, 240–250 (2022).

22. Karczewski, K. J. et al. Systematic single-variant and gene-based association testing of thousands of phenotypes in 394,841 UK Biobank exomes. Cell Genomics 2, 100168 (2022).

23. Madsen, B. E. & Browning, S. R. A Groupwise Association Test for Rare Mutations Using a Weighted Sum Statistic. PLOS Genet. 5, e1000384 (2009).

24. Li, B. & Leal, S. M. Methods for Detecting Associations with Rare Variants for Common Diseases: Application to Analysis of Sequence Data. Am. J. Hum. Genet. 83, 311–321 (2008).

25. Brandes, N., Linial, N. & Linial, M. PWAS: proteome-wide association study—linking genes and phenotypes by functional variation in proteins. Genome Biol. 21, 173 (2020).

26. Wu, M. C. et al. Rare-Variant Association Testing for Sequencing Data with the Sequence Kernel Association Test. Am. J. Hum. Genet. 89, 82–93 (2011).

27. Lee, S. et al. Optimal Unified Approach for Rare-Variant Association Testing with Application to Small-Sample Case-Control Whole-Exome Sequencing Studies. Am. J. Hum. Genet. 91, 224–237 (2012).

28. Li, X. et al. Dynamic incorporation of multiple in silico functional annotations empowers rare variant association analysis of large whole-genome sequencing studies at scale. Nat. Genet. 52, 969–983 (2020).

29. Monti, R. et al. Identifying interpretable gene-biomarker associations with functionally informed kernel-based tests in 190,000 exomes. Nat. Commun. 13, 5332 (2022).

30. Sun, J., Zheng, Y. & Hsu, L. A Unified Mixed-Effects Model for Rare-Variant Association in Sequencing Studies. Genet. Epidemiol. 37, 334–344 (2013).

31. Susak, H. et al. Efficient and flexible Integration of variant characteristics in rare variant association studies using integrated nested Laplace approximation. PLOS Comput. Biol. 17, e1007784 (2021).

32. Zaheer, M. et al. Deep Sets. Adv. Neural Inf. Process. Syst. 30, (2017).

33. Privé, F. et al. Portability of 245 polygenic scores when derived from the UK Biobank and applied to 9 ancestry groups from the same cohort. Am. J. Hum. Genet. 109, 12–23 (2022).

34. Szustakowski, J. D. et al. Advancing human genetics research and drug discovery through exome sequencing of the UK Biobank. Nat. Genet. 53, 942–948 (2021).

35. McLaren, W. et al. The Ensembl Variant Effect Predictor. Genome Biol. 17, 122 (2016).

36. González-Pérez, A. & López-Bigas, N. Improving the Assessment of the Outcome of Nonsynonymous SNVs with a Consensus Deleteriousness Score, Condel. Am. J. Hum. Genet. 88, 440–449 (2011).

37. The ENCODE Project Consortium. An integrated encyclopedia of DNA elements in the human genome. Nature 489, 57–74 (2012).

38. Roadmap Epigenomics Consortium et al. Integrative analysis of 111 reference human epigenomes. Nature 518, 317–330 (2015).

39. Ghanbari, M. & Ohler, U. Deep neural networks for interpreting RNA-binding protein target preferences. Genome Res. 30, 214–226 (2020).

40. Zhou, W. et al. SAIGE-GENE+ improves the efficiency and accuracy of set-based rare variant association tests. Nat. Genet. 54, 1466–1469 (2022).

