## Supplementary Methods for "Integration of variant annotations using deep set networks boosts rare variant association genetics"

October 26, 2023

**8 Supplementary algorithms** **14**

### 1 DeepRVAT model

#### 1.1 Method overview

**Model** The DeepRVAT gene impairment module is trained as part of an end-to-end multi-phenotype prediction model. This model combines gene scores from the gene- and phenotype-agnostic impairment module using phenotype-specific weights to predict one or more phenotypes. During training, we restrict the genes used for phenotype prediction to a set of *seed genes* with known associations to the phenotypes of interest.

**Applications** Subsequently, using the trained DeepRVAT gene impairment module, impairment scores for all protein coding genes can be computed. The derived continuous scores can be leveraged in various downstream tasks, e.g., finding phenotype-associated genes through rare-variant association testing in large cohorts, such as the UK Biobank (UKBB). In addition, DeepRVAT gene impairment scores may be leveraged to augment conventional polygenic risk score (PRS)-based phenotype predictors by including the DeepRVAT scores to capture rare variant effects.

In the remainder of this section, we present a comprehensive description of the DeepRVAT model and how it is trained. Downstream analyses are described in subsequent sections.

#### 1.2 Input and target data

**Variant sets** The input data for DeepRVAT consists of unordered sets of variants. The variant set for individual $i$ and gene $j$ is defined as

$$V_{ij} = \{(a_{kl})_{l=1,\dots,d} \mid \text{variant } k \text{ present in individual } i, \text{ gene } j\}, \tag{1}$$

where $a_{kl}$ is the $l$-th annotation of variant $k$. Thus, $V_{ij}$ is an unordered set of $d$-dimensional vectors, with each vector representing the $d$ annotations of a given variant.

**Targets** As targets, we use a collection of $P$ phenotypes, with the $p$-th phenotype of indivdidual $i$ denoted by $y_i^{(p)}$. We used quantitative (continuous-valued) phenotypes in this study, though we note that an extension to categorical phenotypes is straightforward. DeepRVAT is trained in a multi-task model setup to predict all $P$ phenotypes simultaneously (details below).

#### 1.3 Gene impairment module

The DeepRVAT gene impairment module builds on a set neural network architecture to learn a trait-specific but gene-agnostic scoring function in a data-driven manner [1]. Specifically, the gene impairment module (denoted $\psi$ in what follows) operates on sets of annotated variants and outputs a scalar score.

**Architecture** The variant set is first passed through a learnable submodule $\varphi$, which computes a *variant embedding* $\varphi(x)$ for each $x = (a_{k1}, \dots, a_{kd}) \in V_{ij}$. From the full set of variant embeddings, a fixed aggregation function $f$ computes a *gene embedding*, and then a second learnable submodule $\rho$ computes a scalar *gene impairment score* $\psi(V_{ij})$ from this embedding. In full,

$$\psi(V_{ij}) = \rho\left(f\left(\{\varphi(x)\}_{x \in V_{ij}}\right)\right)$$

Both $\varphi$ and $\rho$ are multi-layer perceptrons (MLPs), i.e., feed-forward neural networks with multiple fully connected layers.

**Aggregation function** The aggregation function $f$ is required to be permutation-invariant, from which permutation invariance of the full phenotype prediction network follows. We also require that $f$ produces outputs at a fixed dimension, independent of the number of elements in the variant set $V_{ij}$, since the MLP $\rho$ requires fixed-dimensional input. Possible choices for $f$ are the element-wise sum, product, or maximum, with output dimension $d$, or the `top2` function, with output dimension $2d$, containing the two largest values of each annotation.

#### 1.4 Training procedure on seed genes

**Seed gene** Training of the gene impairment module begins by selecting, for each phenotype of interest, a set of *seed genes* from the set of all protein-coding genes. These can be based on prior knowledge or, as we do in this study, on the results of alternative RVAT methods, specifically the "Burden/SKAT combined" method described in Section 2.3.

**End-to-end phenotype prediction model** For a given individual $i$, we form variant sets $V_{ij}$ for all seed genes $j$ and compute the gene impairment score for gene $j$ as $\psi\left(V_{ij}\right)$. Following this, we estimate the $p$-th phenotype $y_i^{(p)}$ for individual $i$ as a linear combination of the gene impairment scores and covariates:

$$\hat{y}_i^{(p)} = \boldsymbol{X}_i^T \boldsymbol{\alpha}^{(p)} + \sum_{j \in S^{(p)}} w_j^{(p)} \psi\left(V_{ij}\right),$$

where $\boldsymbol{X}_i$ is the vector of covariates for individual $i$, $\boldsymbol{\alpha}^{(p)}$ is a learnable vector of weights for the covariates, $S^{(p)}$ is the set of seed genes for phenotype $p$, and $w_j^{(p)}$ is a learnable weight for gene $j$.

**Loss function** We employed a simple multi-task learning objective across phenotypes, with the loss function given by the mean loss across all phenotypes. That is,

$$\mathcal{L}(y_i, \hat{y}_i) = \frac{1}{P} \sum_{p=1}^{P} \mathcal{L}(y_i^{(p)}, \hat{y}_i^{(p)}),$$

where again, $P$ is the total number of phenotypes. The weights $\boldsymbol{\alpha}^{(p)}$ and $w_j^{(p)}$, as well as the parameters of $\psi$, are learned via backpropagation and minibatch gradient descent.

**Parameter sharing** The parameters of $\psi$ are shared across all variants, genes, and phenotypes, while $\boldsymbol{\alpha}^{(p)}$ and $w_j^{(p)}$ are phenotype-specific. For the downstream analyses we describe in this work, the covariate and gene weights $\boldsymbol{\alpha}^{(p)}$ and $w_j^{(p)}$ are not needed – only the optimized gene impairment module $\psi^*$ is used.

#### 1.5 Model implementation and hyperparameters

We provide here the specific modeling setup as considered for the results reported in the manuscript. We note that the results obtained using DeepRVAT are robust to moderate adjustments to its hyperparameters; the choices presented here give optimal results while retaining high computational efficiency.

**Software versions** All DeepRVAT models were implemented in PyTorch v1.9.1 and PyTorch-Lightning v1.5.10.

**Training-validation split** A data point for the DeepRVAT multi-phenotype model was given by an individual-phenotype pair. That is, the target was that individuals value for phenotype $p$ and the input was that individual's sets of annotated rare variants, one set for each seed gene specified for phenotype $p$. Prior to training, data points were shuffled, and a validation set consisting of 20% of individuals was selected at random for each phenotype. The validation set was randomly chosen on a per-phenotype basis, meaning it could differ across phenotypes.

**Architecture hyperparameters** For the variant embedding $\varphi$, we used a two-layer MLP with width 20. We used an MLP with two hidden layers of width 10 for $\rho$. In both networks, leaky ReLUs with negative slope 0.01 were used as the activation functions.

**Training hyperparamters** During training, we used the mean-squared-error (MSE) loss and the AdamW optimizer [2] with learning rate 0.0001. The batch size was 1024. During training, the MSE loss was monitored every epoch on the validation set, and the training checkpoint with lowest validation MSE loss was retained. Training proceeded for a minimum of 50 and a maximum of 1,000 epochs, with early stopping implemented by the PyTorch-Lighting `EarlyStopping` callback with metric the validation MSE, `patience` 3 and `min_delta` $10^{-5}$.

#### 1.6 Practical considerations for input data

**Variant and annotation data** Input data preparation first requires to compute all annotation scores for all variants present in the analyzed cohort if they lie in the genomic regions to be considered (e.g., all protein-coding genes). Next, using the cohort genotype data together with genome annotations, the variant set for each individual–gene combination must be determined to obtain matrices of `annotations x variants`. These matrices for each individual and each gene are then combined to obtain the final input tensor. In our practical implementation of DeepRVAT, we achieved the greatest computational efficiency by forming a padded tensor with dimensions of `individuals x genes x annotations x variants`, where, in a given minibatch, the final dimension is padded to the largest number of variants in any individual/gene combination. For model training, the input tensor only comprises the seed genes, with one input tensor used per phenotype.

**Phenotypes and covariates** Besides the variant information, DeepRVAT training and association testing requires the true phenotype values (`individuals x phenotypes`) and the covariates (`individuals x covariates`) to be considered (e.g., age, sex, genetic principal components).

### 2 Relationship with prior work and comparison partners

Broadly, rare variant association testing methods can be categorized into burden tests (also called collapsing tests) and variance-component tests. To increase power, both kinds of tests require a grouping of variants into variant sets (e.g., genes), and incorporate filtering to include only putative causal variants (e.g., pLOF and/or missense variants). They also require applying a predetermined weighting function to individual variants.

#### 2.1 Principles of burden tests and SKAT

The most widely-used methods remain conventional burden tests and the sequence kernel association test (SKAT). Although these models do not conduct any specific modeling to handle more complex annotations, they remain canonical baselines and underpin many more advanced choices [3, 4].

**Burden tests** Burden tests [5] collapse variant weights by, e.g., sum or a binary indicator (presence/absence of a qualifying variant), into a single score per sample and variant set. This score is then tested for association with traits. This simple form of burden test, which is also the most commonly applied, assumes all filtered variants are causal and have the same direction of effect.

**Variance component tests** Variance component tests such as SKAT [6] treat variant effects as random effects and assume that the effects in a given variant set are normally distributed around zero. They test against the null hypothesis that the distribution of effects is a delta distribution at zero. Thus, these tests can handle situations where variants of opposite effects and non-causal variants are present.

**SKAT and SKAT-O** SKAT is a specific type of variance component test for association, utilizing a kernel matrix to estimate the similarity between individuals based on their genotype. More specifically, the kernel in SKAT is given by $K = GWWG^T$, where $G$ is the $n \times p$ genotype matrix and $W = diag(w_1, \ldots, w_p)$ is a specified vector of variant weights. An extension of SKAT, SKAT-O [7], interpolates between SKAT and burden tests, based on the observation that a burden test can be viewed as a special type of kernel test. Annotations enter both SKAT and burden tests as part of the preprocessing of the input data. Variants are typically filtered by allele frequency and pLOF or missense annotations (see also [8, 9] for guidelines and examples). This can comprise simply including all variants of one type (e.g., missense or pLOF [3]), or additional annotations indicating the degree or probability of deleteriousness may be taken into account (e.g., [4]). Also, instead of commonly used variant weighting based on variant MAF only, single or combinations of annotation scores might be used to weight variant in both burden test and SKAT [10].

#### 2.2 Existing methods that handle more complex variant annotations

Building on SKAT and burden tests, there has been a history of model refinements and advances to allow for making use of more complex annotations. In the following, we review the most relevant developments and their relationship to DeepRVAT (see also Supp. Table. 1).

**Hierarchical mixed-effect models** An early example of a method that utilizes functional annotations in a data-driven manner is MiST [11], which introduces a hierarchical mixed-effects model for RVAT. Later, BATI [12] has been proposed as a Bayesian generalization of this model, which uses Integrated Nested Laplace Approximation to improve robustness. While being capable of considering different variant characteristics jointly, both models are linear and learn the relevance of annotations indivdually for each gene, which in practice limits the number and complexity of annotations that can be considered.

**Meta-models** Another emerging group of methods, which we classify here as *meta-models*, perform post-hoc meta-analysis of multiple tests, which can comprise different variant filters (e.g., pLOF or missense variants), test types (e.g., burden test or SKAT), and/or weightings based on single annotation scores or groups of related annotations [13, 14, 15, 10].

**Kernel methods for multiple annotations** For example, Konigorski et al. [15] and Monti et al. [10] proposed linear mixed-effect models that can learn variant effects based on variant weights or on annotations directly. They also developed kernels which allow for user-specified mappings of variants into a feature space, with variant effects modeled linearly based on their representations in feature space. Roughly, then, variant effects are modeled based on their similarity in feature space. Practically, multiple tests for different types of variants are used, each of which defines a weighting or variant similarity kernel based on a set of related annotations.

**STAAR** The STAAR method [14] takes a different perspective on combining multiple annotations into a single test, by first testing for association between each single annotation and a trait, then using statistical methods (namely, the Cauchy combination test [16, 17]) to combine the p-values from these individual tests into a single score. Additionally, the authors use PCA to reduce the total number of annotations tested and the empirical cumulative distribution of annotations to estimate the probability of a variant being causal.

**Limitations of existing methods** While both STAAR and the methods of Konigorski et al. and Monti et al. have been shown to be scalable to handle biobank-scale datasets, these meta-models are limited to test individual variant annotations or groups of related annotations in isolation rather than jointly. They are also not suitable for computing a single gene score that can be used in related tasks, such as phenotype prediction.

**Novel characteristics of DeepRVAT** These existing RVAT methods and our method share some conceptual advances over burden tests and SKAT, including data adaptivity, the usage of functional annotations—including those derived from predictions of deep neural networks—and the mapping of variants into feature space. However, to the best of our knowledge, our approach is the first to learn, in a data-driven manner, nonlinear feature mappings from sequencing studies, and also the first to directly incorporate deep neural networks into an RVAT framework. Furthermore, DeepRVAT contrasts with these methods in that it learns from signal across multiple genes during training time, applying these jointly learned variant representations to other genes in the association testing phase. Finally, DeepRVAT is a single model that is directly applicable to phenotype prediction, while methods based on meta-models require a scheme to combine predictions across multiple models, and have not been considered for this task.

#### 2.3 Implementation of comparison partners

We compared DeepRVAT to burden tests, SKAT, STAAR, and Monti et al. In the following, we provide details on the implementation of these.

##### 2.3.1 Burden tests and SKAT

**Test types, variant annotation** We implemented burden and SKAT tests following [3] using the score test from the `SEAK` package[1] (v0.4.3) [15, 10]. For quantitative (resp. binary) phenotypes, a null model was computed using the `ScoreTestNoK` (resp. `ScoreTestLogit`) class, followed by calling the `pv_alt_model()` method to compute a p-value.

All combinations of burden and SKAT tests restricted to either pLOF or missense variants were carried out, giving four method/variant type combinations. The pLOF variant category comprised all variants annotated as stop gained, start lost, splice donor, splice acceptor, stop lost or frameshift by Ensembl Variant Effect Predictor (VEP) [18, 4]. Annotation of missense variants was also carried out using VEP. Each method/variant type combination was carried out for all protein-coding genes. To reduce the data sparsity due to ultra-rare variants in SKAT tests, variants with a with minor allele count (MAC) $\leq 10$ were collapsed and then tested together with all other variants with MAC $> 10$ as described by [19]. Due to computational constraints, we skipped genes with over 5000 markers, impacting only one gene (Titin) for missense variant tests.

**Variant weights** The weight for each variant $v_j$ was $w_j = \mathrm{Beta}(\mathrm{MAF}(v_j); 1, 25)$ where $\mathrm{Beta}(\cdot; 1, 25)$ denotes the Beta density function with parameters $(1, 25)$, which upweights rarer variants.

**Combination test, multiple testing correction** In addition to the four individual tests, we created a combination test (*Burden/SKAT combined*) using the full set of p-values from all four individual tests. These were then corrected for multiple testing using the Benjamini-Hochberg procedure. The resulting significant gene-trait associations from this combined test were subsequently used as seed genes for training DeepRVAT (see Sec. 1 below).

##### 2.3.2 STAAR

**Software** STAAR tests were implemented in R using the `STAAR` package provided by the authors[2] and following the vignette provided in the package, as well as the procedures in the original publication [14].

**Variant annotation** STAAR requires annotations of variants, and to insure optimal comparability with DeepRVAT, the same annotations as described below in Sec. 6 were used. As required for the STAAR procedure, each annotation $a_{jk}$ for variant $j$ was PHRED-scaled according to the formula

$$a_{jk}^{\mathrm{PHRED}} = -10 \log_{10}(1 - q_{jk}),$$

where $q_{jk}$ represents the quantile of $a_{jk}$ when considering the distribution of annotation $k$ across all variants in the dataset.

**Variant groups, multiple testing correction** Following [14], STAAR p-values were computed for five variant groups, namely (1) putative loss-of-function (stop gain, stop loss and splice), (2) missense, (3) disruptive missense, (4) putative loss-of-function and disruptive missense, and (5) synonymous variants. We defined disruptive missense variants to be those that were predicted to be both "deleterious" by SIFT [20] and "probably damaging" by PolyPhen2 [21]. This yielded five p-values per gene, which were again corrected for multiple testing using the Benjamini-Hochberg procedure.

[1] `https://github.com/HealthML/seak`
[2] `https://github.com/xihaoli/STAAR`

###### 2.3.3 Monti et al.

We conducted various collapsing and kernel tests following the methodology described by Monti et al.(2022)[10]. We used the same annotations, variant weight thresholds, and variant kernel architectures as outlined in their study. Annotation scores were obtained according to the details provided in Supp. Table 5 and Sec. 6.

**Test types** Specifically, we performed the following tests for different types of variants:

- Protein loss of function: Gene-based collapsing test.
- Missense variants: Weighted gene-based variant collapsing and kernel-based association tests. We used SIFT and PolyPhen2 scores for variant weighting and a local (amino-acid level) collapsing kernel to aggregate variants.
- Splicing: Weighted gene-based variant collapsing and kernel-based association tests. We used SpliceAI delta scores for variant weighting together with a linear kernel.
- RBP-binding: We employed a weighted, kernel-based association test using DeepRiPe predictions for six RBPs (QKI, MBNL1, TARDBP, ELAVL1, KHDRBS1, and HNRNPD). Following the approach described in Monti et al. (2022), the kernel matrix was generated using the Cholesky decomposition of the element-wise product of $Q \circ R$, where $Q$ represents the similarity of variants based on their six DeepRiPe scores and $R$ represents the similarity based on variant position.

**Combination of p-values** For all the aforementioned tests, we utilized the score test from SEAK. In the case of missense and splicing tests, if either the collapsing or kernel-based association test yielded nominal significance ($p < 0.01$), we performed joint testing with pLOF variants. The p-values from these tests were integrated using the Cauchy combination method, as described in [10]. In total, we obtained six p-values per gene for each phenotype, which were subsequently adjusted for multiple testing using the Benjamini-Hochberg procedure.

###### 2.3.4 Expected allele frequency filtering

Since burden tests, SKAT, and STAAR use variant filters to define qualifying variants (e.g., pLOF, missense, or disruptive missense), we followed the methodology of [3] to improve the reliability of the tests. Specifically, we restricted testing to genes that passed an expected allele frequency (EAF) filter of at least 50. The EAF is defined as $\sum_{i,j} g_{ij}$, where $g_{ij} \in \{0, 1, 2\}$ is the genotype of sample $i$ and variant $j$, and the sum is taken over all qualifying variants $j$ in the gene in question.

### 3 Association testing with DeepRVAT

#### 3.1 Rare variant association tests

**DeepRVAT impairment testing** Once a gene impairment module $\psi^*$ is trained as in the previous section, gene impairment scores $h_{ij} = \psi^* (V_{ij})$ are computed for each sample $i$ and each gene $j$ to be tested. Using the score test from `SEAK` as in 2.3.1, we then compute an effect size and p-value for the assocation of $h_{ij}$ with the phenotype,

$$\hat{y}_i^{(p)} = \boldsymbol{X}_i \boldsymbol{\gamma}^{(p)} + \beta_j^{(p)} h_{ij}.$$

$\boldsymbol{X}_i$ is the vector of covariates for sample $i$, and $\boldsymbol{\gamma}^{(p)}$ is a learned vector of weights.

**Handling seed genes** When testing for association to phenotypes on which the model was trained, the seed genes used for training must be excluded, since using gene impairment scores from an overfit model may lead to inflated p-values for the genes on which it was fit. Genes not used during model training may be used in association testing and yield calibrated p-values, cf. 3.3.

**EAF filter** Since DeepRVAT considers both high- *and* moderate-impact variants, the EAF filter as described in 2.3 did not have any effect on the set of genes included in association testing.

**Multiple testing correction** If seed genes were discovered using alternative methods on the same dataset used for training and association testing with DeepRVAT, then the p-values for all tests (conventional methods and DeepRVAT) can be combined and multiple testing correction performed on the full set of p-values.

#### 3.2 Repetitions and multiple testing correction

**Background** Commonly, in conventional RVATs, multiple association tests for a given gene/phenotype combination are carried out, each using a different scheme for filtering and/or weighting variants (e.g., one set of tests for pLOF variants and one for missense variants [3], see also [22, 4]). Despite the resultant increase in the multiple testing correction burden, this can increase yield by capturing more diverse aspects of variant effects on the phenotype.

**Stochastic repeats** Since each training run of DeepRVAT begins with stochastically initialized weights, we achieved a similar effect by simply repeating the DeepRVAT training and association testing procedure (Supp. Fig. 3.7). This yielded multiple models $\psi^*$, each of which had an associated uncorrected p-value for every gene.

##### 3.3 Controlling for overfitting

In the design of the DeepRVAT training and association method, we endeavored to control for the potential of false positives caused by overfitting of the Deep Set network. Here, we present an evidence-backed theoretical argument for how this control is achieved, as well as corroborating tests on real and simulated data.

**Theoretical arguments** Recall that the DeepRVAT network is trained on a set of seed genes. These are intentionally not used during the association testing phase, since it is possible that the network has overfit on them during training—meaning that any p-values for associations computed on these genes are not reliable. In constrast, we expect that a network that has overfit will show poor generalization and produce inaccurate phenotype predictions when using non-seed genes. Thus, for an overfit model, we expect that a non-significant p-value should be produced for nearly all genes during association testing.

**Gene-gene correlations** The above argument relies on the assumption that, across samples, the gene-gene correlation between annotated variants is low. Otherwise, even a network that overfits but generalizes poorly could produce accurate phenotype predictions on genes that are highly correlated with the seed genes. Biologically, we have no reason to expect that the existence of, say, a pLOF variant in one gene should correlate with the existence of such a variant in another gene in the same individual. Indeed, we find on our the UKBB cohort that this gene-gene correlation between variants is low (Supp. Fig. 3.6). It is, on the whole, even lower than the sample-sample correlation between variant types in a given gene, justifying DeepRVAT's training-association testing split by gene rather than by sample.

**Empirical evidence** Finally, we recall Fig. 2, where DeepRVAT's false discovery rate (FDR) was verified, on the basis of simulations, to be calibrated, and Fig. 3, where we see a high replication rate for DeepRVAT discoveries, implying good generalization. These provide additional lines of evidence, on both simulated and real data, that overfitting is controlled in the method.

##### 3.4 Leveraging pre-trained DeepRVAT models on held-out traits and cohorts

In addition to using DeepRVAT for association testing on traits that have been used during training of the DeepRVAT gene impairment module as described in 3, it may be applied to any trait or cohort not considered during training.

To perform association testing (as in Section 3), gene impairment scores $h_{ij}$ are calculated for all sample/gene pairs of interest using the pre-trained DeepRVAT gene impairment modules across all repeats, as detailed in Section 3.2. These modules require variant annotation vectors $(a_{kl})$ with the same $d$ annotations used during DeepRVAT gene impairment module training. Once these computed scores are computed for the desired gene/individual pairs, they can be used to discover associations with quantitative or binary traits of interest, resulting in one p-value per tested trait and DeepRVAT repeat.

#### 4 Phenotype prediction using DeepRVAT and alternative rare variant scores

##### 4.1 Problem statement

**Tasks** Here, we combine conventional common variant polygenic risk scores (PRS) with rare variant gene burdens, comparing the performance of DeepRVAT gene burden scores with alternative scores. Two separate problems were addressed: the prediction of raw phenotype values, and prediction of high-risk individuals. We separately assessed two binary high-risk individual stratification tasks: (1) lowest 0.01-th quantile (ranked by phenotype value) vs. rest, and (2) highest 0.01-th quantile vs. rest.

**Cross-validation** In each case, individuals were randomly split into five folds for cross-validation (CV).

**PRS computation** Common PRS variants and effect sizes were all obtained from the Polygenic Score (PGS) Catalog [23] using the study from [24]. The catalog numbers of each common variant PRS are listed in Supp. Table 3.

##### 4.2 Alternative burdens

**Gene discovery** On each CV fold, the training individuals were used for gene discovery using the method "Burden/SKAT combined" described in Sec. 2.3. Retaining associations at an FDR $< 0.05$ resulted in a set of genes $G_b^{(p)}$ for phenotype $p$ to use in the baseline prediction models.

**Burden scores** Subsequently, for each sample $i$ (training and validation), each gene $j \in G_b^{(p)}$, and each annotation $k$ considered, burdens $s_{ij}^k$ were computed by taking the maximum of annotation $l$, i.e.,

$$s_{ij}^l = \max_{k \in W_{ij}} a_{kl}, \tag{2}$$

where $W_{ij}$ denotes the set of all variants $k$ in sample $i$ and gene $j$, and $a_{kl}$ is the $l$-th annotation of variant $k$.

#### 4.3 DeepRVAT gene impairment scores

**Gene discovery** For each CV fold, the set of genes $G_b^{(p)}$ – for the 21 phenotypes $p$ indicated in Supp. Table 3 – from the previous section were utilized as seed genes to train a DeepRVAT gene impairment module $\psi_r^*$ as described in Sec. 1, with $r = 1, \ldots, R$ indexing the repeats. Subsequently, the trained DeepRVAT gene impairment modules were used for association testing on all 33 traits of interest[3] as in Sec. 3, which yielded a set of trait-associated genes $G_d^{(p)}$ with an FDR $< 0.05$ when correcting for the full set of DeepRVAT repeats and "Burden/SKAT combined" tests. Both training and association testing were carried out only on the training samples for that fold, leaving the validation samples held out.

**Gene impairment scores** Finally, DeepRVAT gene impairment scores $\psi_r^*(V_{ij})$ were computed on the variant set $V_{ij}$ (cf. (1)) for each sample $i$ and $j \in G_d^{(p)}$ using the $R$ trained gene impairment modules from the previous paragraph.

#### 4.4 Phenotype predictor training and evaluation

For simplicity, we describe models for predicting raw phenotype values; prediction of extreme values is analogous, with logistic regression on the binary target replacing linear regression.

**Baseline** As a baseline phenotype predictor, we consider a regression model where the explanatory variables comprise covariates (age, sex, the first 20 genetic PCs) and the common variant PRS score:

$$\hat{y}_i^{(p)} = \boldsymbol{\alpha}^T \boldsymbol{X}_i + \beta_c^{(p)} c_i^{(p)},$$

where $c_i^{(p)}$ is the common variant PRS score of sample $i$ for phenotype $p$ and, as above, $\boldsymbol{X}_i$ is the vector of covariates for sample $i$. The weights learned during regression were $\boldsymbol{\alpha}$ and $\beta_c^{(p)}$.

**Extension with rare variants** To incorporate the effects of rare variants into the phenotype predictors, we extended the common variant PRS models by the rare burden scores of significant genes, with models incorporating DeepRVAT or alternative burdens given respectively by

$$\hat{y}_i^{(p)} = \boldsymbol{\alpha}^T \boldsymbol{X}_i + \beta_c^{(p)} c_i^{(p)} + \sum_{j \in G_d^{(p)}} \beta_j^{(p)} \psi_r^*(V_{ij}),$$
$$\hat{y}_i^{(p)} = \boldsymbol{\alpha}^T \boldsymbol{X}_i + \beta_c^{(p)} c_i^{(p)} + \sum_{j \in G_b^{(p)}} \beta_j^{(p)} s_{ij}^p.$$

The difference lies in whether DeepRVAT or alternative burdens are used, and additionally the burdens and learned gene weights $\beta_j^{(p)}$ range over either the "Burden/SKAT combined" gene set $G_b^{(p)}$ or the DeepRVAT gene set $G_d^{(p)}$. The effect of gene set choice is further assessed in Supp. Fig. 4.1. Additionally, since DeepRVAT training is repeated $R$ times, we fit $R$ separate regression models, using burdens computed from the respective gene impairment modules, and average the results.

**Model fitting** The linear and logistic regression models were fit in `R` v4.2.0 using the functions `lm` and `glm` (resp.) from the `stats` package using the family `binomial()` for logistic regression models, and otherwise retaining the default parameters. For logistic regression, the minority class samples were upsampled to match the number of samples in the majority class.

**Evaluation** All models were fit on the training samples for the respective CV fold. The trained models were used to generate predictions on the validation samples of that fold. Combining these predictions across all CV folds gave exactly one prediction per sample. All evaluations were carried out on this set of predictions.

[3]Waist-to-Hip Ratio was excluded, since no PRS scores were available for this phenotype

##### 4.5 Phenotype predictor assessment

**Prediction accuracy** To start, we assessed the performance of the phenotype predictors using two metrics: coefficient of determination ($R^2$) for the linear models and area under the precision-recall curve ($AUPRC$) for the logistic models. We compared the performance of two phenotype predictors using only the common variant PRS or using both the common PRS and rare variant burdens. Next, we calculated the relative improvement of the model that leverages rare variant burdens compared to the common PRS-only model as

$$\text{Relative}\Delta M = \frac{M_{rare} - M_{PRS_{only}}}{M_{PRS_{only}}},$$

where $M$ denotes $AUPRC$ or $R^2$, respectively. We compared the relative improvement of the rare variant model using DeepRVAT gene impairment scores to predictors using alternative rare variant burdens by a one-sided Wilcoxon test.

**Additional evaluations** The next evaluation focused on individuals with strong deviations between the rare variant and common variant PRS predictor. For each phenotype and individual, we calculated the absolute difference between the predicted phenotype values obtained from a linear model using either the common variant PRS alone or the the common variant PRS together with the rare variant burdens. Subsequently, we ranked individuals based on the magnitude of this difference. At each rank, we determined the count of individuals exhibiting outlier phenotypes, specifically those falling within the top or bottom 1% of the phenotypic distribution. Finally, we tested the enrichment of phenotype predictor outliers in individuals with extreme phenotypes. Across a range of z-score phenotype outlier cutoffs, we identified individuals above the phenotypic cutoff and determined the proportion of these individuals with a predicted phenotype value exceeding the 99% quantile. Enrichment scores were scaled relative to the baseline population (z-score = 0) and compared to the common PRS-only model.

#### 5 Simulations

##### 5.1 Genotype data

To compare power and type I error rates of DeepRVAT to conventional RVAT methods in simulation studies, we used semi-synthetic datasets generated based on real genotypes and annotations from the UK Biobank WES interim 200k release (more details in Sec. 6 below). To minimize confounding due to population structure, we restricted to 167,245 individuals of Caucasian genetic ethnicity as determined by an analysis of genetic principal components [25].

##### 5.2 Overview of phenotype simulation

Here, we give an overview of the phenotype simulation scheme. For the precise algorithm and the parameters used see Algorithm 1 and Supp. Table 2.

**Simulated causal genes** We first sampled 100 genes and designated these as *simulated causal genes*. Gene sampling was restricted such that all sampled genes passed an expected allele frequency (EAF) filter of at least 50 for missense variants, and that 50% passed this filter for pLOF variants.

**Variant impact score** Next, variants in the simulated causal genes with MAF below the specified threshold were assigned an impact score based on selected variant annotations, namely variant MAF, 11 binary annotations (VEP consequences) and 5 continuous scores (SIFT, PolyPhen2, Condel, PrimateAI, CADD, AbSplice). (For more details on variant annotations, see Sec. 6.) Binary annotations were assigned a weight inversely proportional to their frequency, thereby reflecting the assumption that infrequent annotations, for example loss-of-function, have a larger impact. Continuous annotations were equally weighted. The variance explained by each variant score component (continuous annotations, binary annotations, MAF and variant score noise) is flexibly tunable.[4]

**Simulated causal variants, and gene burdens** Based on the impact score (with noise added to introduce stochasticity), simulated causal variants were selected, such that a specified proportion of all variants was designated to be causal. Using these, gene burden scores were computed for each sample and each simulated causal gene by first aggregating variants individually for each annotation, followed by computing the weighted sum across the aggregated variant annotation scores. The genetic component of the simulated phenotype was then obtained as the sum of the gene burdens.

[4]We note that since the annotations are not independent of one another, what we describe as the "variance explained" by each component cannot precisely be understood as such—for example, the sum of the "variance explained" by each component will be greater than the total variance explained by the annotations. Nevertheless, for simplicity, we will slightly abuse nomenclature by continuing to refer to this as "variance explained."

**Simulated phenotypes** Finally, to obtain the simulated phenotype values, we randomly sampled covariates and noise and rescaled each phenotype component (covariates, genetics, and noise) such that it explained the specified variance.

**Allele frequency spectrum** In Fig. 2b, we varied the effect size explained by variants with larger allele frequencies. To achieve this, we modified the weight assigned to binary annotations by employing weights defined as $q^{-1/x}$ for variant weighting, where $q$ denotes the allele frequency (AF) and $x$ takes on the values of 1, 3, and 10. As $x$ increases, a greater number of variants with higher allele frequencies were selected as causal. The cumulative effect size explained by variants in each AF bin was determined as follows. Initially, we assigned a per-variant effect size to all simulated causal variants, which was computed by multiplying the simulated variant weight by the AF. Next, we calculated the cumulative effect size explained by each AF bin by summing the per-variant effect sizes within that particular bin and dividing this sum by the total summed effect sizes of variants from all AF bins. The cumulative effects explained by each AF bin were averaged across multiple simulation repeats.

#### 5.3 Assessment of simulation results

**Calibration** To assess statistical calibration, phenotypic data was simulated from the null by sampling from a standard normal distribution, i.e., with no simulated causal genes or variants.

**Power and type I error rates** For all other simulation experiments, statistical power and type I error rates were assessed by comparison of discoveries (FDR $< 0.05$ as determined by the Benjamini-Hochberg procedure) to the simulated causal genes. For the rank-based evaluation (Supp. Fig. 2.3b), we ranked gene-trait associations by their $p$-value as computed by DeepRVAT and the alternative methods. At each rank, we determined the cardinality of the intersection between the gene-trait associations at or below that rank and the set of simulated causal genes, and finally averaged the cardinality across simulation repeats.

### 6 Application to UK Biobank WES data

#### 6.1 Sequencing data preprocessing and quality control

**Exome sequencing** Whole-exome sequencing (+100 bp overhang) was performed on 200,633 participants from the UK Biobank [26], for which the methods have been described for the earlier release of data from approximately 50,000 individuals [27]. While larger WES cohorts from UK Biobank are now available, we restricted to the smaller cohort for this initial methodological study in order to enable the assessment of replication rates as described below.

**Variant data and QC** Variant calling data was downloaded from the UK Biobank as project-level VCF (pVCF) files. Since the dataset had not been subjected to variant- or sample-level filtering prior to release by the UK Biobank, we applied additional quality control(QC) following [27]. All filtering steps were performed using bcftools v1.10.2 [28]. Briefly, we required a minimum read depth of 7 for SNPs and 10 for indels. After read-depth filtering, only variant sites with at least one homozygous variant genotype or where at least one sample per site had an allelic balance ratio greater than 15% for SNPs and 10% for indels were retained. Indels were left-aligned and normalized, and multi-allelic variants were represented as multiple bi-allelic variants. Finally, we removed duplicate variants and filtered for variants where the fraction of missing genotypes was $< 10\%$ and the Hardy-Weinberg equilibrium p-value was $> 10^{-15}$. In total, our filtering criteria removed 5,336,543 out of 17,981,684 initial variants (29.67%). After excluding sex chromosomal variants, we ended up with 12,417,590 variants. Additionally, we filtered out individuals with $> 10\%$ missing genotype rate. In order to minimize any confounding due to population structure and help ensure any discoveries were based on the direct signal from rare variants, we retained only individuals of Caucasian ancestry. This filtering resulted in a final dataset of 167,245 individuals.

#### 6.2 Custom sparse genotype data format

**Raw data** The storage size of the pVCF files for the UK Biobank WES dataset was 8.4 TB, which posed a major challenge for data processing. Furthermore, after QC, we required the data to be stored in a format that allowed for fast, repeated loading over multiple epochs of DeepRVAT model training. To meet our needs, we constructed a custom sparse genotype data format as follows.

**Genotype extraction** After using the bcftools `norm` function for normalization and left alignment, we obtained BCF output files. Next, we extracted triplets of variant metadata, sample ID, and genotype from the BCF files, where the genotype was encoded as 1 (heterozygous) or 2 (homozygous-alternative), thereby generating a set of gzipped TSV files of 71 GB total, with one line for every variant present in a sample. Homozygous-reference genotypes were ignored for the purposes of these files. Following this, each unique variant was assigned an integer ID.

**Custom HDF5 format** As the last step, we created our custom sparse dataset in Hierarchical Data Format 5 (HDF5 v1.10.6). Genotype data was encoded as two equal-sized matrices, a variant and a genotype matrix, with the rows corresponding to individuals and the number of columns equal to the maximum number of variants found in any sample (60,247). Each row of the variant matrix provided the IDs of variants present in the corresponding individual, while the row of the genotype matrix provided the corresponding genotypes (1 or 2). Unnecessary elements at the end of each row were padded with -1. Sample IDs corresponding to the rows of the matrices were stored as an additional sample vector. Details on variants such as position and chromosome, as well as reference and alternative allele, were provided as a variants dataframe in Apache Parquet format. In total, the HDF5 dataset had a storage size of approximately 100 GB.

#### 6.3 Covariates and variant annotation

**Covariates** We retrieved genetic sex, sample age and the first 20 genetic principle components (PCs) directly from UK Biobank (Supp. Table 4). All of these covariates were included in association testing and when training DeepRVAT.

**Variant-to-gene assignments** Variants were assigned to genes using those genes and exons marked as golden in the merged Ensembl/HAVANA genome annotations (GENCODE release 38). We assigned a variant to a gene if it was located at most 300 bp from an exon of that gene.

**Annotations** The full collection of variant annotations used and their sources is provided in Supp. Table 5. Here, we give details on processing for those annotations which were not used directly in the form output by the source.

**MAF** MAF values for variants were first replaced with the maximum of the MAF in the UK Biobank cohort and in gnomAD release 3.0 (non-Finnish European population). Following [5], each MAF $p_j$ was then transformed according to the formula $[p_j(1-p_j)]^{-\frac{1}{2}}$ for use in modeling.

**VEP consequences** We used 11 moderate- and high-impact consequences from VEP. These were encoded for each variant as multi-hot vectors, with a 1 in the corresponding column indicating that a consequence was predicted, and 0 if not.

**DeepSEA** DeepSEA predicts 919 different predicted variant effects on transcription factor binding, DNase I sensitivities, and histone marks in various cell types [29]. To improve model fitting and avoid overfitting, we performed principal components (PC) analysis and restricted to the first 6 PCs, explaining approximately 58% of variance.

**SpliceAI** SpliceAI provides four "delta scores" indicating a variant's predicted effect on cryptic splicing (acceptor gain, acceptor loss, donor gain, and donor loss) [30]. We computed the maximum of these four scores and used it as a single annotation.

**AbSplice** AbSplice-DNA predicts variant effects on aberrant splicing across 49 human tissues [31]. We computed the maximum predicted effect across tissues and used this as a single annotation.

**DeepRiPe** DeepRiPe characterizes in vivo RNA binding protein (RBP) binding preferences [32]. As in [10], we predicted effects of genetic variants on the binding of six RBPs over three cell lines using the pre-trained models from [33].

**Annotation vectors** Given the full set of annotations and the transformations described above, vectors representing variants had 33 dimensions in total.

**MAF thresholds** For DeepRVAT training (see below), we used variants with MAF $< 1\%$. For association testing with all methods, we designated rare variants as having MAF $< 0.1\%$. Additionally, for both training and association testing, we restricted to variants with PHRED-scaled CADD value $> 5$.

#### 6.4 Phenotype data

All phenotype data was obtained directly from UK Biobank.

**Quantitative traits** In case multiple instances of the phenotype were available, we chose the instance with the largest number of individuals having a measurement. Following previous work [10], we selected a set of blood biochemistry measurements as representative quantitative phenotypes, while also adding blood count phenotypes. In addition, we included standing height as a well-studied polygenic trait. Phenotype values were quantile transformed to match their empirical distributions to a standard normal distribution. Suppl. Table 3 provides an overview of the full set of quantitative phenotypes and their corresponding data fields in UK Biobank.

**Binary traits** Binary traits were extracted using the definitions from [22, Supp. Table 1]. Phenotype values were set to 1 for an individual if any "Matching Code" was found for the corresponding "Field" in that table. Otherwise they were set to 0. The exception is if the corresponding "Exclude" value was 1: In this case, the phenotype was set to `NA`.

##### 6.5 DeepRVAT training and association testing

The DeepRVAT gene impairment module was trained as described in Sec. 1 using the 21 phenotypes indicated in Supp. Table 3, and variants from all seed genes discovered by the baseline methods (Sec. 2.3.1).

For the traits used during DeepRVAT training, association testing was carried out on all genes not used as seed genes following the procedure in Sec. 3 using six DeepRVAT model repeats (using more yielded diminishing returns, Supp. Fig. 3.8) and correcting for multiple testing considering all repeat models as outlined in Sec. 3.2. For binary and quantitative traits not used during gene impairment module training, association testing was carried out on all genes, again using six DeepRVAT model repeats. For consistency with the case of testing using training phenotypes, we also applied the seed gene discovery method to new traits and included the resulting p-values in the multiple testing correction. In Figure 3, we report only the number of unique genes that were significant at a false-discovery rate (FDR) $< 0.05$.

##### 6.6 Assessment of replication

**Reference discoveries** We began by collecting, for all phenotypes used in this study, discovered gene-trait associations from two studies [3, 4] on larger WES cohorts from UK Biobank (454,787 and 394,841 individuals, respectively). We counted as a discovery any association that was considered significant according to the methodology of the study. (For [3], we included tests using missense or pLOF burdens and genes that met the EAF filter as described in 2.3.4.) We call this set of discoveries

$$D^{(C)} = \{(j, p) \mid \text{ gene } j \text{ significantly associated to phenotype } p\}$$

**Calculation of replication** Next, for each method $m$, we collected gene-trait associations $(j, p)$ for all genes $g$ and phenotypes $p$, and ranked them by p-value, resulting in a list $(j_1, p_1), \ldots, (j_N, p_N)$. The discoveries of rank $m$ or less are then $D_m = \{(j_q, p_q) \mid 1 \leq q \leq m\}$, and the replication rate at rank $m$ is defined as $|D_m \cap D^{(C)}|$.

##### 6.7 Assessment of calibration

We repeated the same procedure for DeepRVAT training and association testing as above, but using phenotype values that were permuted across individuals. To restrict the analysis to an evaluation of DeepRVAT, excluding contributions from the seed gene discovery methods, the seed genes used were those discovered on true phenotype values. Additionally, Q-Q plots, genomic inflation factors, and discovery counts in Fig. 1c and Supp. Fig. 3.4 refer to results derived from DeepRVAT impairment scores only, not including the seed gene discovery methods.

#### 7 Evaluation of feature importance

Despite their enormous success in improving predictive performance for various tasks, deep learning models remain challenging to interpret as to explain the model's decision on certain outcomes, and thus are commonly referred to as black-box models. In genomics, in-silico mutagenesis is one way of explaining the impact of the input perturbations on the model outcome through forward propagation. However such experiments are computationally expensive, thus encouraging the use of alternative approaches such as DeepLIFT [34] and GradCAM [35]. For a given input, these back-propagation based solutions compute the contribution of each feature by backpropagating through the network from the corresponding prediction. In order to explain the importance of the variant annotations used in DeepRVAT, we conducted separate analyses for quantitative and binary annotations to provide comparisons within each of these types.

##### 7.1 Quantitative annotations

**SHAP** We employed SHAP (SHapley Additive exPlanations), a game-theory based method that assigns an importance score for each feature (in our case, annotation) based on the change in the expected model prediction for each corresponding model output [36]. In particular, we used SHAP DeepExplainer (v0.41.0), which integrates the DeepLIFT algorithm to approximate SHAP values, to compute importance scores for all variant annotations that were used to train DeepRVAT for phenotype prediction.

**Subsampling** For each repeat used in Sec. 6 and associated trained gene impairment module, the SHAP DeepExplainer was trained on 3,000 individuals from the training set and the importance of the variant annotations were explained on 1,000 individuals from the validation set. (The training and validation sets were the same as those used in Sec. 6.) Subsampling was necessary due to computational constraints.

**Importance scores** SHAP values produced by the explainer have the same shape as the input data, which is `individuals x genes x annotations x variants`. We aggregated the absolute SHAP values on individual, gene and variant levels to obtain an importance vector with the size of `annotations` by computing the average. Finally, the SHAP value vectors from each repeat were averaged to obtain aggregated importance scores.

**Robustness of subsampling** In order to illustrate the robustness of subsampling for feature importance explanations, we repeated the above procedure 15 times, each time randomly selecting different subsets of individuals from the training and validation sets. The annotations highlighted as important by the SHAP DeepExplainer were robust to the changes in the training and validation individuals. The final feature importance analyses were based on the aggregated SHAP values of the 15 different subsamplings.

#### 7.2 Binary annotations

The impact of binary variant effect annotations was measured through in-silico mutagenesis experiments where only training (seed) genes were considered. For each binary annotation $\hat{l}$, we first created a filtered subset of individuals ($S_{\hat{l}}$) where each individual and gene pair has at least one annotated variant $v_k = (a_{kl})$ where $a_{k\hat{l}} = 1$ (wild type) in a given phenotype. A complementary mutant subset ($M_{\hat{l}}$) was created from $S_{\hat{l}}$ in which the corresponding annotation of the variants ($a_{k\hat{l}}$) were in-silico mutated to $a_{k\hat{l}} := 0$. The wild type ($S_{\hat{l}}$) and mutant ($M_{\hat{l}}$) individual subsets were separately scored using the DeepRVAT gene impairment module. The absolute difference between the gene impairment scores (aggregated over individuals and genes) was considered as the impact of the annotation of interest. Finally, a relative importance score for each annotation was calculated where the annotation with the highest absolute difference value was set to the relative importance value of 1. The relative importance of other annotations were scaled accordingly by dividing their difference scores by the maximum difference score value $A$, i.e., ($a_{\hat{l}}/A$).

#### 7.3 Variant annotation groups

**Groups** Among the variant annotations that are utilized to train DeepRVAT, some have similar functionalities and are highly correlated (Supp. Fig. 3.1). Therefore, in order to increase the interpretability of the feature importance analysis, we grouped certain annotations. Since UKBB MAF, CADD raw and DeepSEA principal components (PC1-PC6) were highlighted as relatively important compared to the rest of the continuous annotations by the SHAP explainer, we treated each of these annotations as a group in itself. The remaining annotations were assigned to a group (see (Supp. Table 5).

**DeepSEA** We also assigned human-readable labels to first 6 DeepSEA principal components (PC1-PC6) that were used as annotations, based on the absolute values of the loadings (Supp. Fig. 3.2) belonging to 919 epigenetic and regulatory genomic tracks. The principal components were named after the feature(s) within the top-ten highest loading values (Supp. Fig. 3.2). For instance, we referred to DeepSEA PC2 as CTCF, since the five out of ten highest ranking loadings belonged to CTCF-related tracks.

### 8 Supplementary algorithms

**Parameters**

- Maximum MAF $MAF_{max}$
- 100 randomly selected simulated causal genes ($g_k$) for $k = 1, ..., 100$
- Variance explained by each variant score component: Minor allele frequency ($V_{MAF}$), binary annotations ($V_{binary}$), continuous annotations ($V_{cont}$), variant score noise ($V_{vnoise}$)
- Proportion of causal variants $p_{causal}$ (across all simulated causal genes), $\#vars\_total$ the total number of variants with MAF $< MAF_{max}$ across all simulated causal genes
- Variance explained by each phenotype component: Genetic component ($V_{gen}$), Non-genetic components: Noise ($V_{pnoise}$), covariates ($V_{cov}$).

**assert** $V_{MAF} + V_{binary} + V_{cont} + V_{vnoise} = 1$
**assert** $V_{gen} + V_{pnoise} + V_{cov} = 1$

**Input data**

- **Genotypes** $G_{ij}$ for sample $i$, $(i = 1, ..., n)$ and variant $j$, $(j = 1, ..., J)$ variants with MAF $< MAF_{max}$
- **Annotation scores** $a_{jl}$ for $(l = 1, ..., L)$ annotations: Minor allele frequency, $A_{binary}$: 11 binary annotations (VEP consequences) $A_{cont}$: 5 continuous annotations scores (SIFT, PolyPhen2, Condel, PrimateAI, CADD, AbSplice DNA score)

**Simulate data**

1. Determine explained variance $V_l$ for each annotation:
   - Binary annotations: $V_l = V_{binary} * 1/f_l \quad l \in A_{binary}$, where $f_l$ is inverse proportional to the frequency of binary annotation $l$ and $\sum f_l = 1$
   - Continuous annotations: $V_l = V_{cont} * 1/\#cont\_annotations \quad l \in A_{cont}$
2. Rescale annotations $A_l$ such that the variance explained by each annotation is $V_l$
$$A_l^* = w_l * A_l$$
, where $w_l$ scales $A_l$ such that $Var(A_l^*) = V_l$
3. Compute variant scores S as
$$S = \sum_{l=1}^{L} A_l^*$$
4. Rank variants according to their score $s_j$ and select the top $p_{causal} \cdot \#vars\_total$ as causal variants
5. Compute gene score $b_{k,i}$ for all samples $i = 1, ..., n$ and all simulated causal genes $k = 1, ..., K$:
$$b_{k,i} = \sum_{l=1,...,L} w_l \sum_{j=1,...,J} a_{jl} \cdot g_{ij}, \quad j \in \text{causal\_variants}$$
6. Aggregate gene burdens to obtain genetic effects $Y^{gen}$
$$y_i^{gen} = \sum_k b_{k,i}$$
7. Sample noise and covariate terms
$$\text{Noise: } Y^{pnoise} \sim N(0,1)$$
$$\text{Covariates: } Y^{cov_1} \sim N(0,1), Y^{cov_2} \sim Bernoulli(0.5)$$
8. Calculate scaling factors $w_{gen}$, $w_{pnoise}$, $w_{cov}$ for the phenotype components ($Y^{gen}$, $\epsilon$, $X$) such that each phenotype component explains the desired proportion $V_{gen}$, $V_{pnoise}$, $V_{cov}$ of the phenotypic variance
9. Compute phenotype values as the sum of the scaled phenotype components
$$Y = w_{gen} \cdot Y^{gen} + w_{cov} \cdot (Y^{cov_1} + Y^{cov_2}) + w_{pnoise} \cdot Y^{pnoise}$$

**Algorithm 1:** Simulation algorithm used to generate simulated phenotype data from UK Biobank WES data.

### References

[1] Manzil Zaheer et al. “Deep Sets”. *Advances in Neural Information Processing Systems* 30 (2017).

[2] Ilya Loshchilov and Frank Hutter. “Decoupled weight decay regularization”. *arXiv preprint arXiv:1711.05101* (2017).

[3] Konrad J. Karczewski et al. “Systematic single-variant and gene-based association testing of thousands of phenotypes in 394,841 UK Biobank exomes”. *Cell Genomics* 2.9 (Sept. 2022).

[4] Joshua D. Backman et al. “Exome sequencing and analysis of 454,787 UK Biobank participants”. *Nature* 599.7886 (Nov. 25, 2021).

[5] Bo Eskerod Madsen and Sharon R. Browning. “A Groupwise Association Test for Rare Mutations Using a Weighted Sum Statistic”. *PLoS Genet* 5.2 (Feb. 13, 2009). Ed. by Nicholas J. Schork.

[6] Iuliana Ionita-Laza et al. “Sequence kernel association tests for the combined effect of rare and common variants”. *American Journal of Human Genetics* 92.6 (June 6, 2013).

[7] Seunggeun Lee et al. “Optimal Unified Approach for Rare-Variant Association Testing with Application to Small-Sample Case-Control Whole-Exome Sequencing Studies”. *The American Journal of Human Genetics* 91.2 (Aug. 2012).

[8] Gundula Povysil et al. “Rare-variant collapsing analyses for complex traits: guidelines and applications”. en. *Nature Reviews Genetics* 20.12 (Dec. 2019).

[9] Seunggeung Lee et al. “Rare-Variant Association Analysis: Study Designs and Statistical Tests”. *The American Journal of Human Genetics* 95.1 (July 2014).

[10] Remo Monti et al. “Identifying interpretable gene-biomarker associations with functionally informed kernel-based tests in 190,000 exomes”. *Nat Commun* 13.1 (Sept. 10, 2022).

[11] Jianping Sun, Yingye Zheng, and Li Hsu. “A Unified Mixed-Effects Model for Rare-Variant Association in Sequencing Studies”. *Genet. Epidemiol.* 37.4 (May 2013).

[12] Hana Susak et al. “Efficient and flexible Integration of variant characteristics in rare variant association studies using integrated nested Laplace approximation”. *PLoS Comput Biol* 17.2 (Feb. 19, 2021). Ed. by Yue Li.

[13] Zihuai He et al. “Unified Sequence-Based Association Tests Allowing for Multiple Functional Annotations and Meta-analysis of Noncoding Variation in Metabochip Data”. *The American Journal of Human Genetics* 101.3 (Sept. 2017).

[14] Xihao Li et al. “Dynamic incorporation of multiple in silico functional annotations empowers rare variant association analysis of large whole-genome sequencing studies at scale”. *Nat Genet* 52.9 (Sept. 2020).

[15] Stefan Konigorski, Shahryar Khorasani, and Christoph Lippert. “Integrating omics and MRI data with kernel-based tests and CNNs to identify rare genetic markers for Alzheimer’s disease”. *arXiv:1812.00448 [cs, q-bio, stat]* (Mar. 5, 2019). arXiv: 1812.00448.

[16] Yaowu Liu and Jun Xie. “Cauchy Combination Test: A Powerful Test With Analytic $p$ -Value Calculation Under Arbitrary Dependency Structures”. *Journal of the American Statistical Association* 115.529 (Jan. 2, 2020).

[17] Yaowu Liu et al. “ACAT: A Fast and Powerful p Value Combination Method for Rare-Variant Analysis in Sequencing Studies”. *The American Journal of Human Genetics* 104.3 (Mar. 2019).

[18] William McLaren et al. “The Ensembl Variant Effect Predictor”. *Genome Biol* 17.1 (Dec. 2016).

[19] Wei Zhou et al. “SAIGE-GENE+ improves the efficiency and accuracy of set-based rare variant association tests”. *Nature genetics* 54.10 (2022).

[20] Prateek Kumar, Steven Henikoff, and Pauline C Ng. “Predicting the effects of coding non-synonymous variants on protein function using the SIFT algorithm”. *Nat Protoc* 4.7 (July 2009).

[21] Ivan A Adzhubei et al. “A method and server for predicting damaging missense mutations”. *Nat Methods* 7.4 (Apr. 2010).

[22] Sean J. Jurgens et al. “Analysis of rare genetic variation underlying cardiometabolic diseases and traits among 200,000 individuals in the UK Biobank”. *Nat Genet* (Feb. 17, 2022).

[23] Samuel A. Lambert et al. “The Polygenic Score Catalog as an open database for reproducibility and systematic evaluation”. en. *Nature Genetics* 53.44 (Apr. 2021).

[24] Florian Privé et al. “Portability of 245 polygenic scores when derived from the UK Biobank and applied to 9 ancestry groups from the same cohort”. en. *The American Journal of Human Genetics* 109.1 (Jan. 2022).

[25] Clare Bycroft et al. “The UK Biobank resource with deep phenotyping and genomic data”. *Nature* 562.7726 (Oct. 2018).

[26] Joseph D. Szustakowski et al. “Advancing human genetics research and drug discovery through exome sequencing of the UK Biobank”. *Nat Genet* 53.7 (July 2021).

[27] Cristopher V. Van Hout et al. “Exome sequencing and characterization of 49,960 individuals in the UK Biobank”. *Nature* 586.7831 (Oct. 29, 2020).

[28] Petr Danecek et al. “Twelve years of SAMtools and BCFtools”. *GigaScience* 10.2 (Jan. 29, 2021).

[29] Jian Zhou and Olga G. Troyanskaya. “Predicting effects of noncoding variants with deep learning–based sequence model”. *Nature Methods* 12.10 (Oct. 2015).

[30] Kishore Jaganathan et al. “Predicting Splicing from Primary Sequence with Deep Learning”. *Cell* 176.3 (Jan. 24, 2019).

[31] Nils Wagner et al. “Aberrant splicing prediction across human tissues”. *Nature Genetics* (2023).

[32] Mahsa Ghanbari and Uwe Ohler. “Deep neural networks for interpreting RNA-binding protein target preferences”. *Genome research* 30.2 (2020).

[33] *DeepRipe*. `https://github.com/ohlerlab/DeepRiPe`. 2022 (Online; accessed July, 2022).

[34] Avanti Shrikumar, Peyton Greenside, and Anshul Kundaje. “Learning important features through propagating activation differences”. In: *International conference on machine learning*. PMLR. 2017.

[35] Ramprasaath R Selvaraju et al. “Grad-cam: Visual explanations from deep networks via gradient-based localization”. In: *Proceedings of the IEEE international conference on computer vision*. 2017.

[36] Scott M Lundberg and Su-In Lee. “A unified approach to interpreting model predictions”. *Advances in neural information processing systems* 30 (2017).
