## Supplementary Information for "Integration of variant annotations using deep set networks boosts rare variant association genetics"

### Supplementary figures

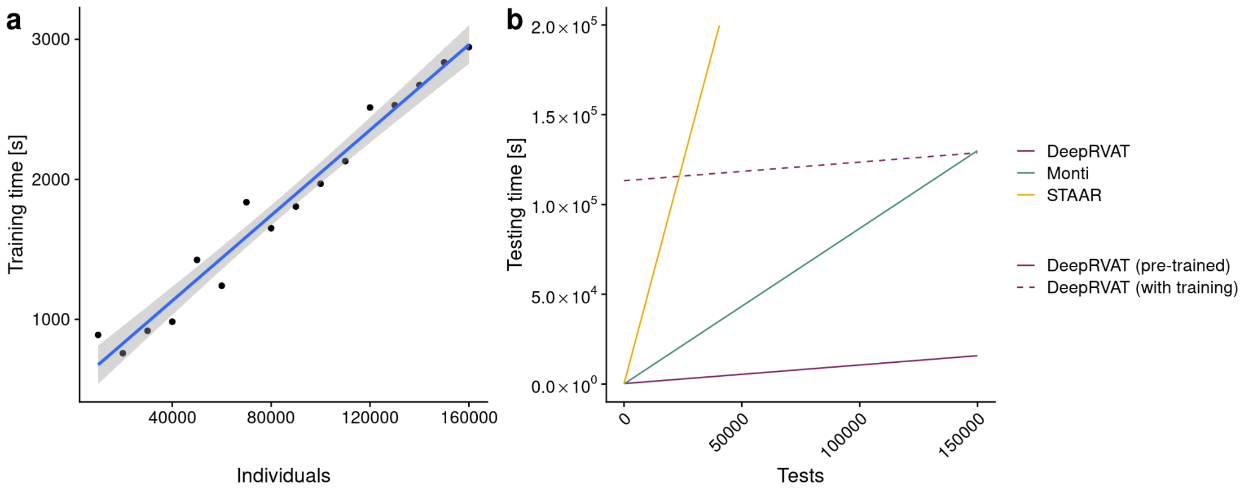


**Supp. Fig. 1.1: Training and association testing times for DeepRVAT and comparison methods.** All timing experiments were conducted on a workstation with an AMD Ryzen Threadripper PRO 5975WX CPU and an NVIDIA RTX 4090 24GB GPU.

(**a**) The UK Biobank WES was downsampled in intervals of 5,000 individuals and DeepRVAT was trained five times for each sample size. The plot indicates the mean training time in seconds, which scales roughly linearly with the sample size.

**(b)** Comparison of association testing times between DeepRVAT and alternative methods. Average compute time was determined by evaluating 1,000 tested genes for a single phenotype, resulting in the average time required to test one gene-phenotype pair. For STAAR and Monti et al.'s, the total test time per gene was calculated by summing the compute times for individual variant filter masks or kernel designs and test types, respectively. The y-intercept of DeepRVAT (with training) represents the time needed for seed gene discovery and model training. The x-axis is truncated at 200,000 seconds.

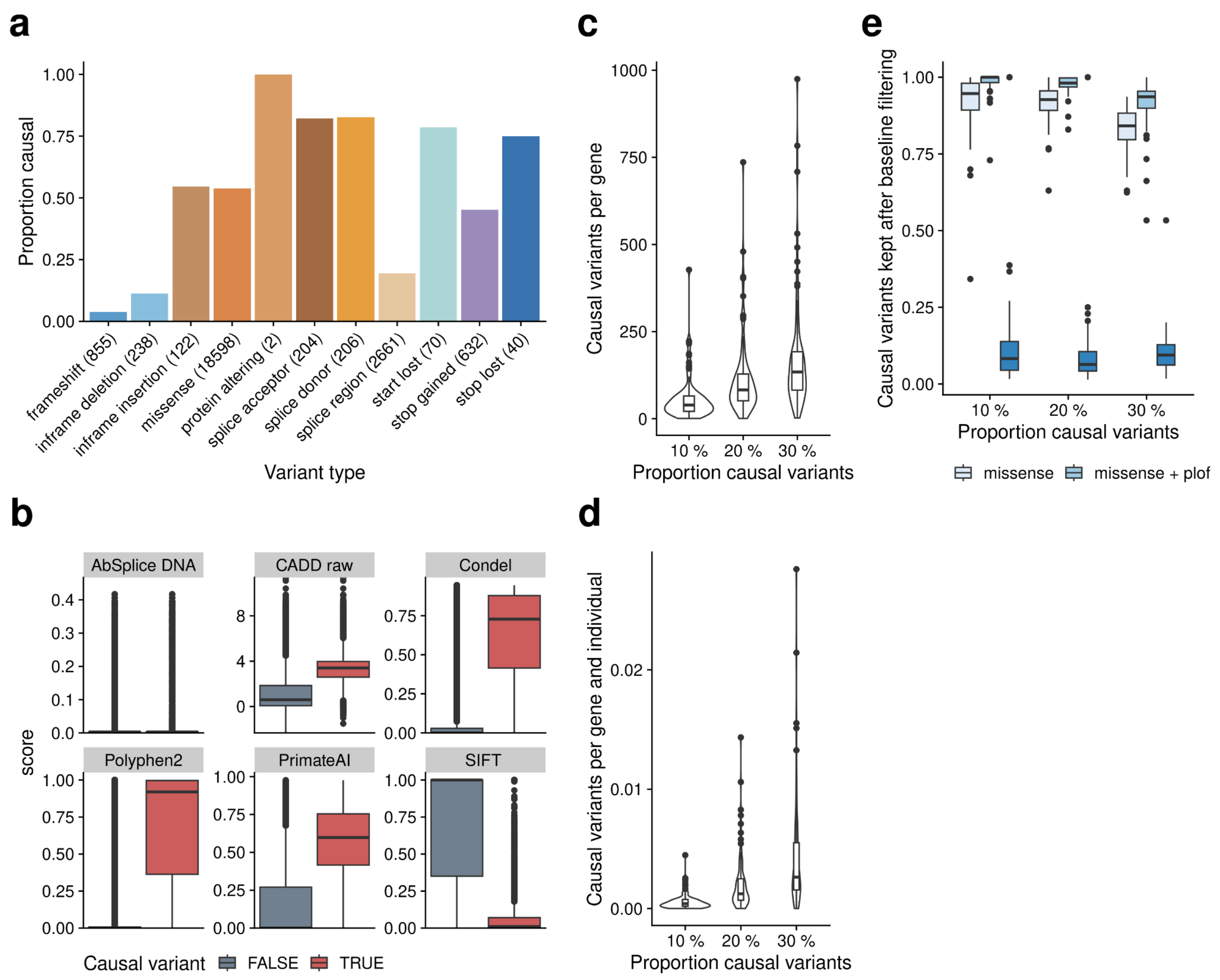


**Supp. Fig. 2.1: Properties of simulated causal variants.** To define causal variants, variants in the simulated causal genes with MAF below the specified threshold (**Supp. Table 4**) were assigned an impact score based on selected variant annotations, namely variant AF, binary and continuous annotations and stochastic noise. Based on the impact score, simulated causal variants were selected such that a specified proportion of all variants was designated to be causal.

including the default value of 20%. Values are averaged across 10 simulation replicates.

**(c)** Number of causal variants for each gene designated to be causal. The observed variation reflects the simulation procedure as high-impact variants are selected irrespective of the gene they belong to.

(**d**) Number of causal variants per gene and individual, averaged across individuals for each gene.

(**e**) Proportion of simulated causal variants that are retained by the missense/pLOF variant filter as used in baseline methods. Across different proportions of causal variants, a high proportion of causal variants passes filtering for the baseline methods when combining missense/pLOF filter masks, which confirms that the baseline methods are, in theory, capable of detecting the simulated associations.

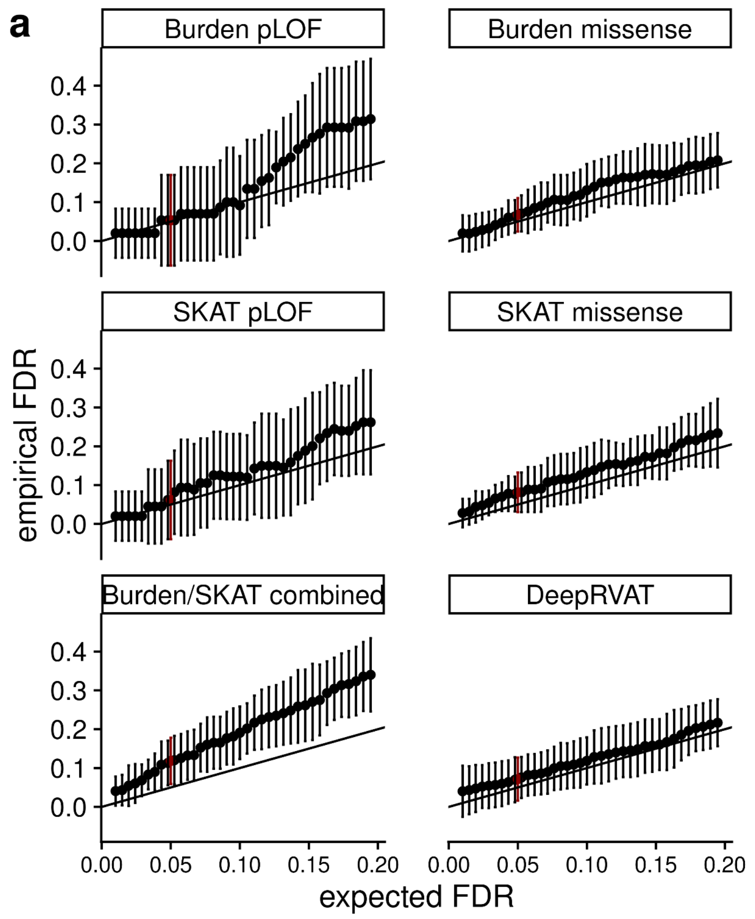


**Supp. Fig. 2.2: Statistical calibration of DeepRVAT.** All results shown are from simulations with the default parameters (**Supplemental Table 4**).

(**a**) Empirical false discovery rate (FDR) versus the expected FDR. Shown are average values and plus or minus one standard deviation for alternative rare variant tests, estimated from 10 simulation replicates.

(**b**) Genomic inflation factor $\lambda_{GC}$ (genomic control) for the same models and data as in **a**. Box plots denote median and lower and upper quartiles across simulation replicates.

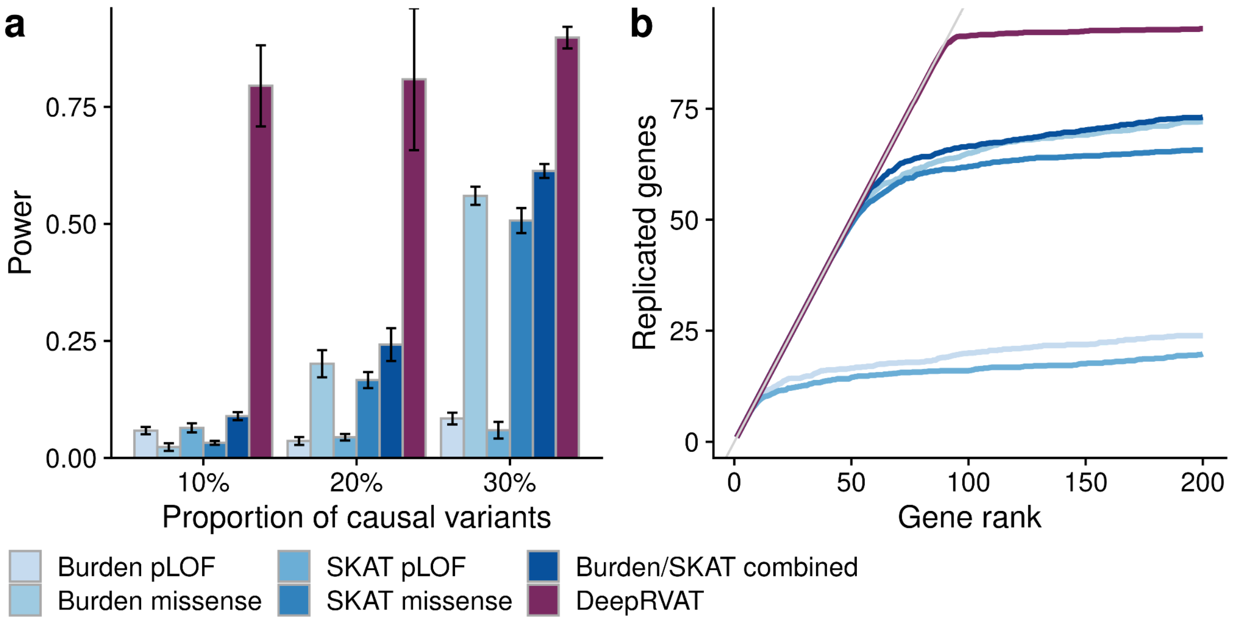


**Supp. Fig. 2.3: Model assessment using simulated data.** Comparison of DeepRVAT using the complete set of annotations versus conventional burden or SKAT tests, using either pLOF or missense annotations, considering different simulation settings.

(**a**) Power of alternative methods for varying proportions of variants simulated to affect the trait. From left to right, 10%, 20%, and 30% of rare variants with MAF < 0.1% in the simulated causal genes were selected as causal variants. Bar height denotes average power across 10 simulated replicates; error bars correspond to plus or minus one standard deviation.

(**b**) Rank-based evaluation. Genes were ranked by p-value and the proportion of simulated causal genes (true positives) at each rank was determined (average over 10 simulation replicates).

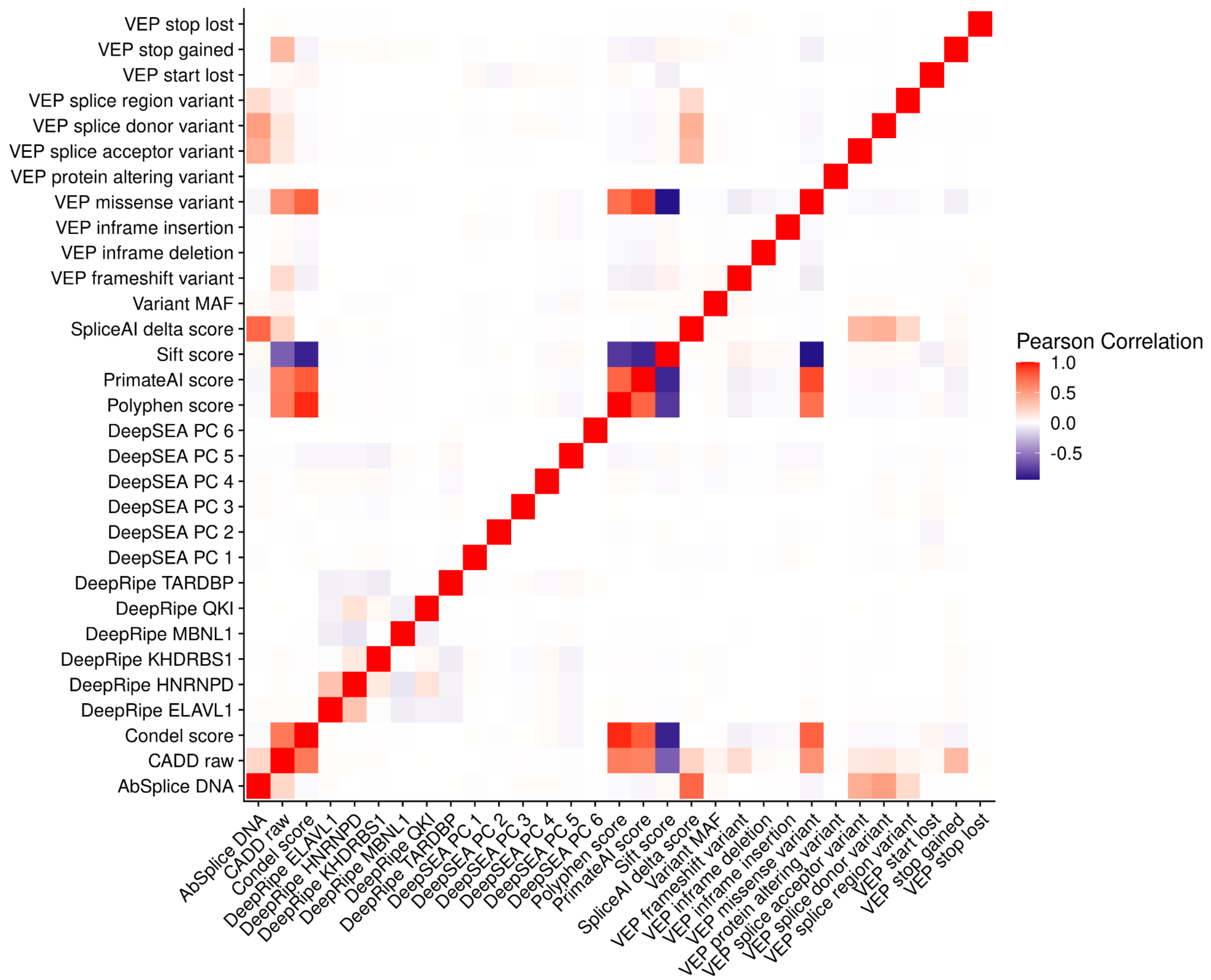


**Supp. Fig. 3.1: Pairwise correlation of functional variant annotations.** Heatmap showing Pearson correlation between 33 individual functional annotations using variants from the UKBB cohort.

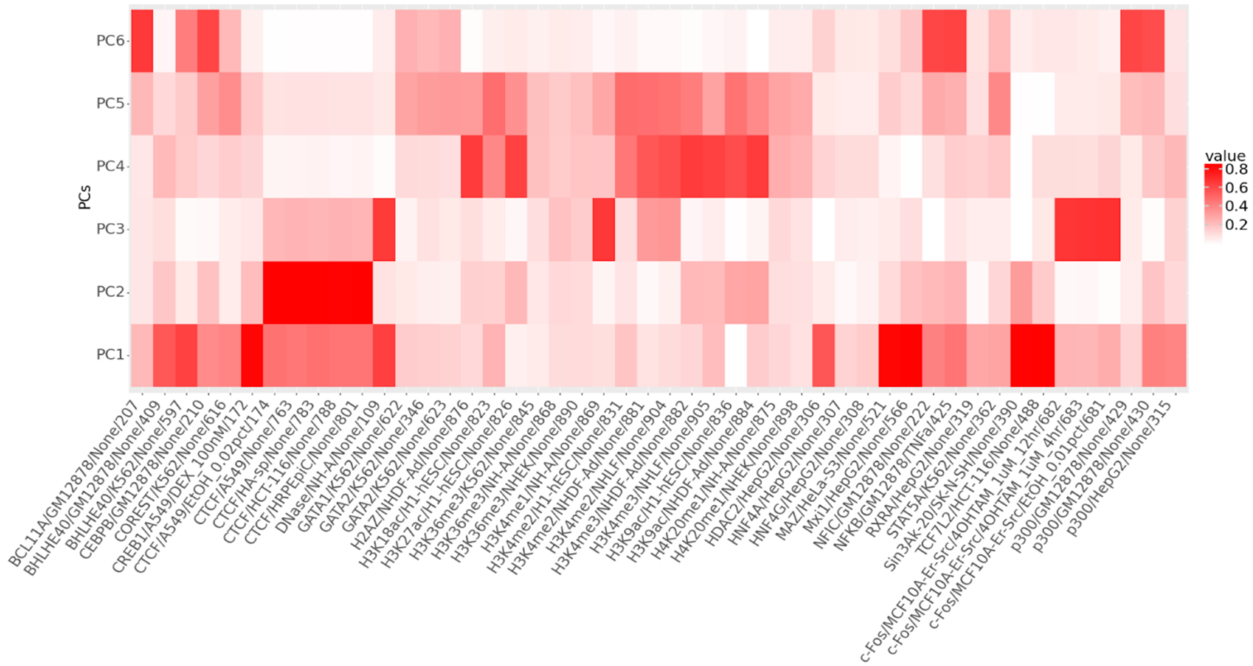


**Supp. Fig. 3.2: Loadings for top six DeepSEA principal components.** The top six DeepSEA principal components were used as annotations within DeepRVAT. Labels indicate feature name, cell-type, treatment-type, and position index among 919 original DeepSEA features.

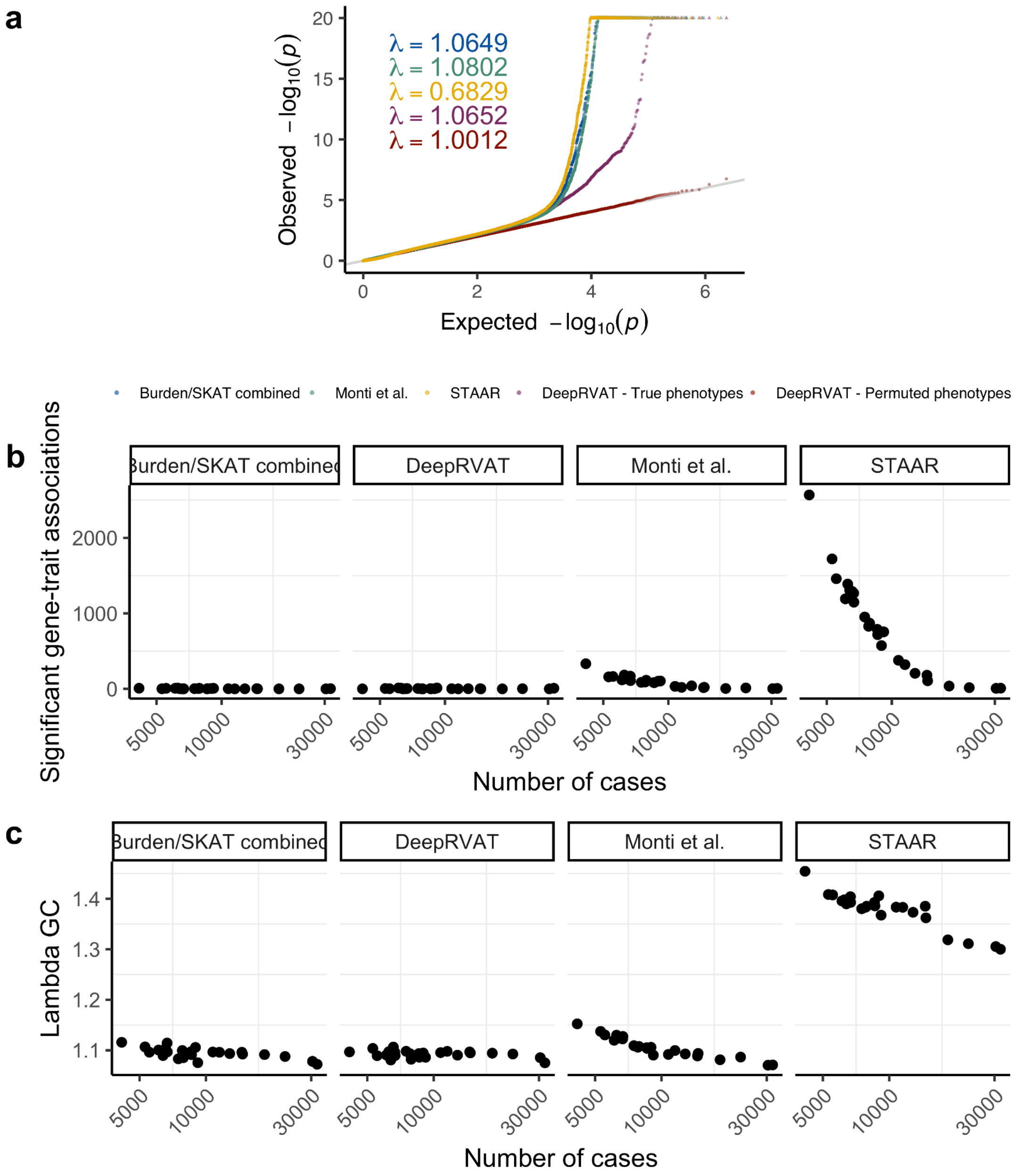


**Supp. Fig. 3.3: Assessment of statistical calibration of alternative methods.**
(**a**) Q-Q plots of the expected vs. the observed *p*-value distribution for alternative methods on quantitative traits. Sown are aggregate results across all methods for Burden/SKAT combined, Monti et al., STAAR and DeepRVAT. DeepRVAT - Permuted phenotypes denotes DeepRVAT applied to permuted phenotype values.
(**b**) Significant gene-trait associations (FDR < 0.05) vs. number of cases for binary traits. Results from Monti and STAAR show a strong relationship between the rarity of cases and the number of discoveries, which reflects the inflated test statistics in **a**.
(**c**) Genomic inflation factor vs. number of cases for binary traits.

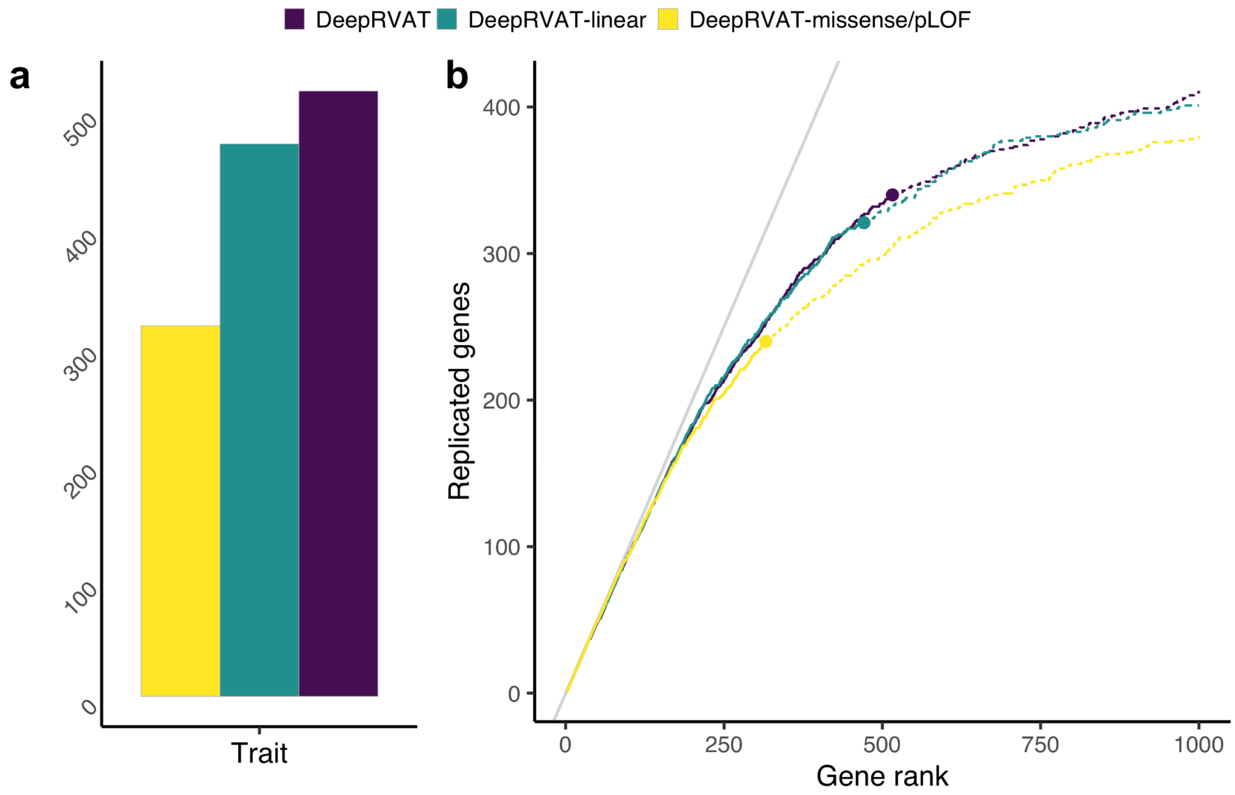


**Supp. Fig. 3.4: Comparison of alternative DeepRVAT configurations.** DeepRVAT was trained using either a reduced set of annotations (MAF, pLOF, and missense status: DeepRVAT-missense/pLOF) or considering a reduced model architecture, incorporating the complete set of variant annotations but employing a fully linear gene impairment module (DeepRVAT-linear). The performance of these alternative DeepRVAT setups was evaluated against the results from the full DeepRVAT model as shown in main text Fig. 3.

(**a**) Cumulative number of gene-trait associations discovered by alternative DeepRVAT setups across 21 traits (FDR<5%).

(**b**) Replication of the cumulative discoveries as in **a** in larger cohorts (defined as in main text Fig. 3b) across all traits. Shown is, for each DeepRVAT setup, the total number of gene-trait associations that were also discovered in the larger cohorts as a function of the rank of their nominal significance. Circles indicate the rank position that corresponds to FDR < 5%.

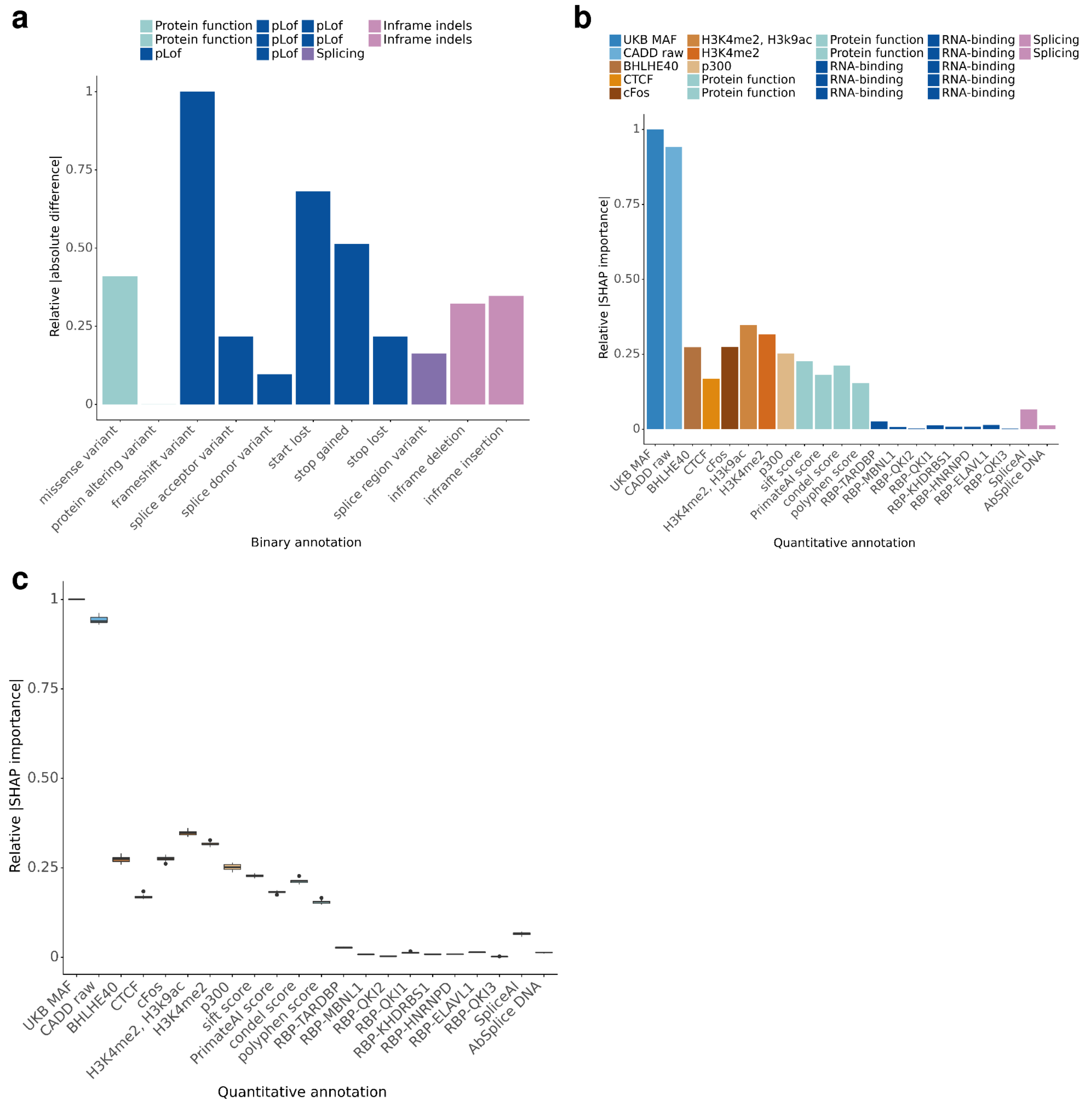


**Supp. Fig. 3.5: Feature importance analysis.** The relative importance scores displayed in plots range between 0-1 and are calculated such that the relative importance of the binary/quantitative annotation with the strongest impact is 1. The annotations are depicted with the same color if they belong to the same annotation group.

**(a)** *In-silico* mutagenesis analysis for binary annotations considered by DeepRVAT. The impact of a given binary annotation was assessed by comparing DeepRVAT predictions using the real annotation vector with a mutated annotation vector, where the effect of the considered binary annotation was masked (set to zero). The absolute difference between the two gene impairment scores, aggregated across all samples and genes, was considered as the impact of the variant of interest.

**(b)** SHAP importance values for quantitative annotations considered by DeepRVAT. In order to assess the contribution of quantitative annotations to the DeepRVAT predictions, we utilized SHAP DeepExplainer. The explainer is trained on 3000 samples from the training set and 1000 samples from the validation set are used to obtain SHAP importance values from the trained explainer. The output of the explainer is the same shape as the input. Therefore the absolute SHAP values are aggregated to obtain annotation-level importance scores. This procedure is repeated 15 times with different training and validation samples. The final scores are computed by the aggregation over 15 different samplings.

**(c)** The relative SHAP importance scores for quantitative annotations are robust across 15 different samplings. The relative SHAP importance scores of quantitative annotations are plotted across 15 samplings, where different training and validation samples are fed to the explainer model.

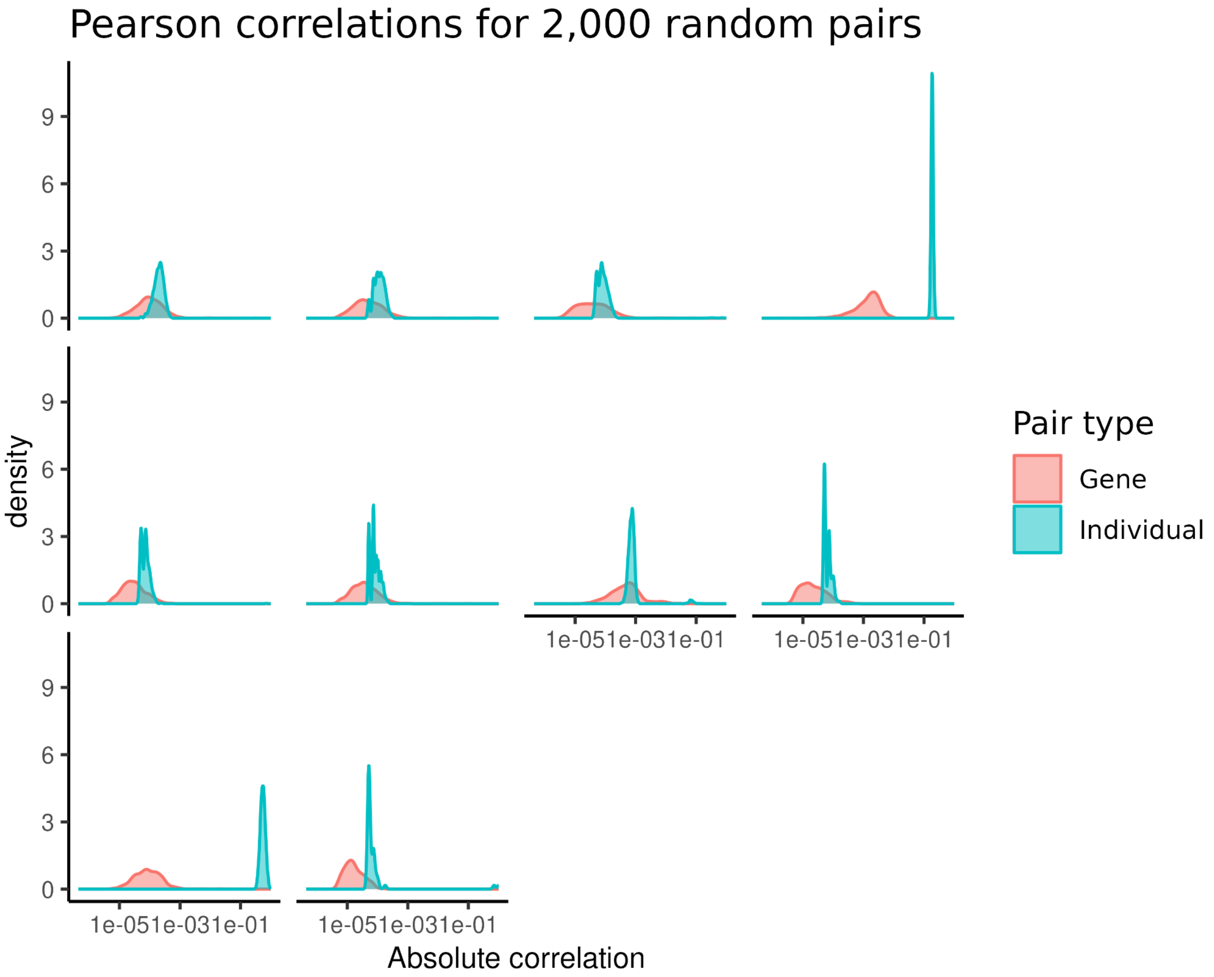


**Supp. Fig. 3.6: Correlation analysis.** For each VEP consequence used as an annotation within DeepRVAT (**Methods**), we computed an individual-gene matrix by summing the number of variants with that consequence in a given gene and individual; data from the UKBB WES dataset. We then randomly sampled 2,000 pairs of individuals (rows) and genes (columns) and computed the correlations of the counts. The plots show the distribution of the absolute values of these correlations.

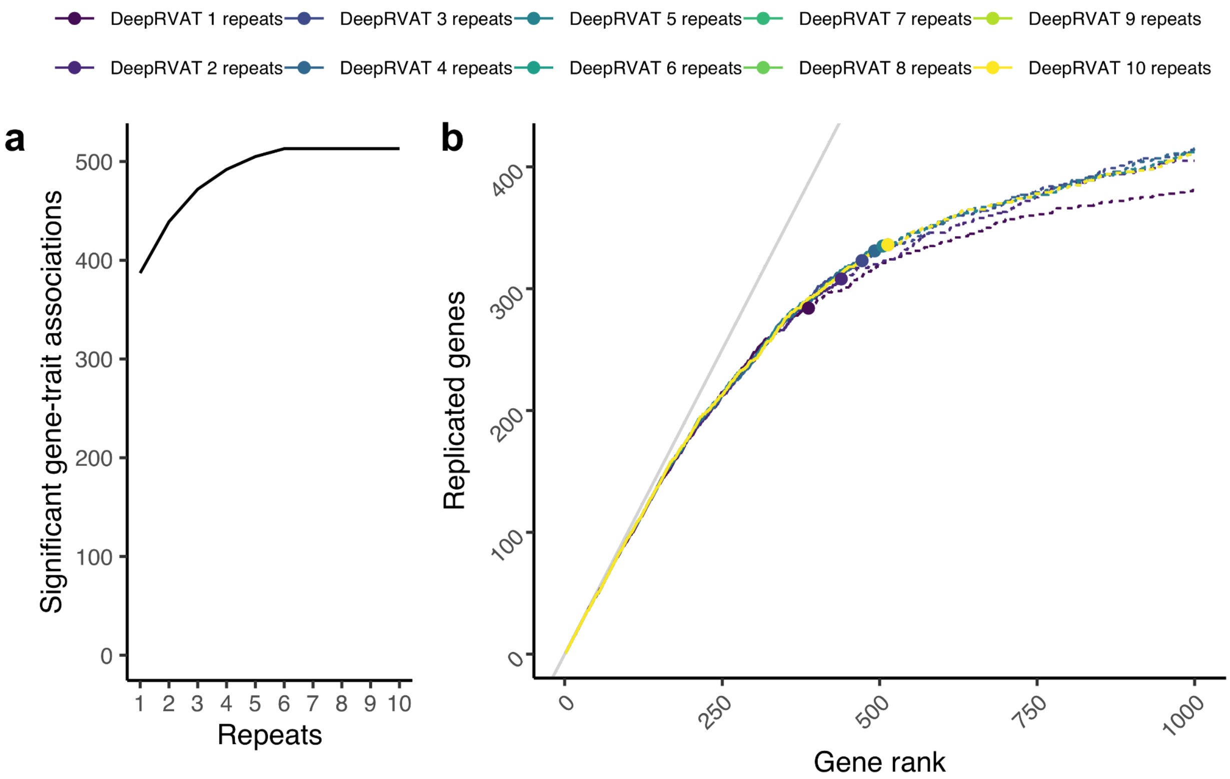


**Supp. Fig. 3.7: Repeat analysis.** Comparison of association testing results on UK Biobank whole-exome sequencing data (as in **Fig. 3**), considering different numbers of repeats.
(**a**) Count of significant gene-trait associations (FDR < 0.05).
(**b**) Replication analysis, analogous to **Fig. 3b**.

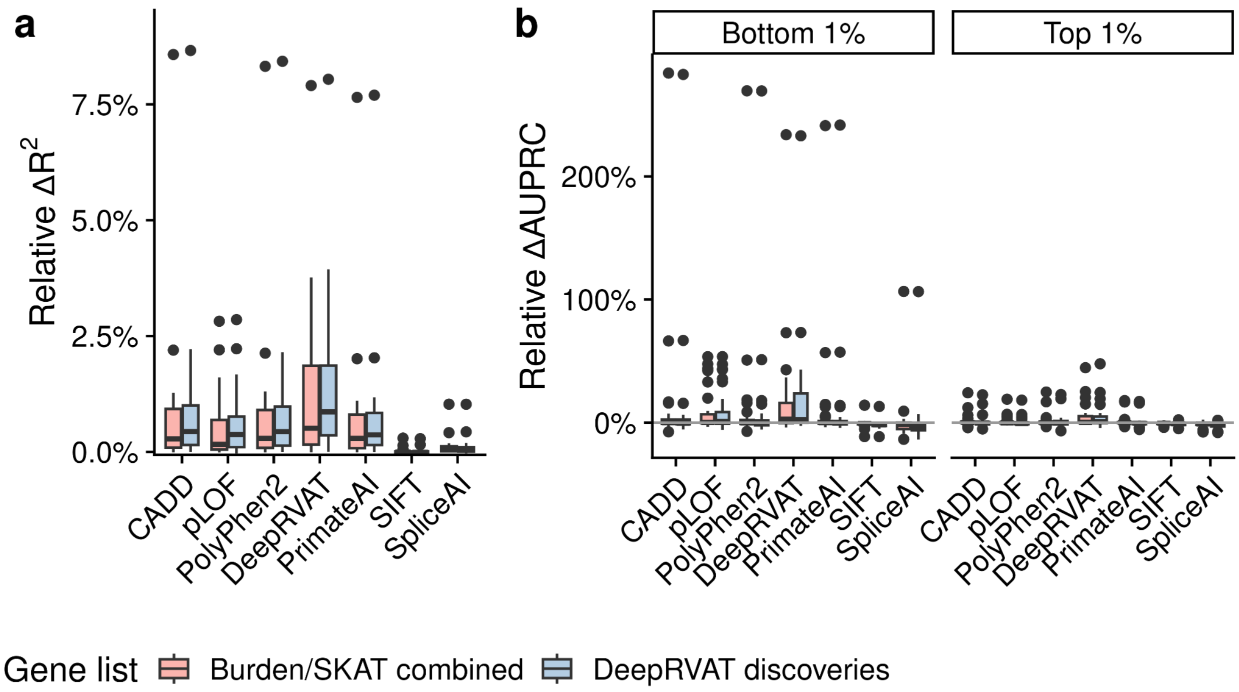


**Supp. Fig. 4.1: Impact of gene list choice on the performance of rare variant phenotype predictors.** For each rare variant burden method, we compared the performance of rare variant phenotype predictors when utilizing rare variant gene burdens derived from genes identified through conventional RVAT methods (Burden/SKAT combined) or DeepRVAT (gene discovery methods applied separately to each fold of the training set) with FDR < 5%.

(**a**) Relative improvement of the prediction performance of a linear regression model that includes rare variant gene burdens versus a model based on common variant PRS only. DeepRVAT gene impairment scores consistently outperform alternative rare variant phenotype predictors, regardless of the gene list choice. Additionally, incorporating genes discovered by DeepRVAT further enhances the performance compared to using genes identified by conventional RVAT methods.

(**b**) Analogous comparison as in **a**, however considering a logistic regression model to stratify individuals in the bottom or top 1% of the phenotypic distribution. Shown are relative differences in the area under the precision-recall curve (AUPRC) between a model that includes rare variant gene burdens versus a PRS-only model.

### Supplementary table captions

**Supplemental Table 1.** Conceptual comparison of DeepRVAT to related methods.

**Supplemental Table 2**. Simulation parameters. For each figure related to the simulation studies, simulation parameters used with the phenotype simulation algorithm as described in **Methods** are listed.

**Supplemental Table 3**. Quantitative and clinical phenotypes from UK Biobank used in this study. Columns: Phenotype: Name of analyzed trait; UKBB Data Field: UK Biobank data field ID corresponding to trait; Number of samples/cases: Number of individuals measured for the trait (quantitative phenotypes) or number of cases (binary phenotypes) and with whole exome sequence data available; PRS ID: PGS catalog id (https://www.pgscatalog.org/) from which variants and effect sizes for the common variant PRS calculation were retrieved; Used for DeepRVAT training: Indicates if the phenotype was used for training DeepRVAT; Trait Group: Trait group the trait was assigned to for plots in Figure 3.

**Supplemental Table 4.** Covariates used as controls association testing as well as DeepRVAT training. Columns: Covariate: Name of covariate; UKB field ID: UK Biobank field ID for covariate.

**Supplemental Table 5**. Variant annotations used within DeepRVAT. The table presents a comprehensive description of each variant annotation utilized, including the source of the annotation score, the assigned group in the feature importance analysis (Supp. Fig. 3.6), and whether the annotation was employed by the method proposed by Monti et al.

**Supplemental Table 6**. Significant gene trait associations discovered by DeepRVAT.

**Supplemental Table 7**. Gene-trait associations discovered in other UK Biobank association studies on larger cohorts. The significant associations, as defined by Backman et al. 2021 or Karczewski et al. 2022 (Genebass), are listed for all traits examined in this paper.
